# Mechanisms of Feature Selectivity and Invariance in Primary Visual Cortex

**DOI:** 10.1101/2020.02.08.940270

**Authors:** Ali Almasi, Hamish Meffin, Shaun L. Cloherty, Yan Wong, Molis Yunzab, Michael R. Ibbotson

## Abstract

Visual object identification requires both selectivity for specific visual features that are important to the object’s identity and invariance to feature manipulations. For example, a hand can be shifted in position, rotated, or contracted but still be recognised as a hand. How are the competing requirements of selectivity and invariance built into the early stages of visual processing? Typically, cells in the primary visual cortex are classified as either simple or complex. They both show selectivity for edge-orientation but complex cells develop invariance to edge position within the receptive field (spatial phase). Using a data-driven model that extracts the spatial structures and nonlinearities associated with neuronal computation, we show that the balance between selectivity and invariance in complex cells is more diverse than thought. Phase invariance is frequently partial, thus retaining sensitivity to brightness polarity, while invariance to orientation and spatial frequency are more extensive than expected. The invariance arises due to two independent factors: (1) the structure and number of filters and (2) the form of nonlinearities that act upon the filter outputs. Both vary more than previously considered, so primary visual cortex forms an elaborate set of generic feature sensitivities, providing the foundation for more sophisticated object processing.

## Introduction

Visual object recognition depends critically on incorporating both selectivity and invariance into the processing of features in the visual input. For example, in inferotemporal cortex neurons are selective to elaborate features that constitute high-level representations of objects (Lehky and Tanaka 2016; Kravitz et al. 2013; DiCarlo et al. 2012; Cadieu et al. 2014; DiCarlo and Cox 2007). At the same time, these neurons are invariant to a range of object transformations such as changes in size, luminance, brightness polarity, orientation, and position, thus allowing for robust object recognition (Kravitz et al. 2013). The visual system combines fine selectivity for particular features, with insensitivity to variations in features that are of little importance to the object’s identity (Lehky and Tanaka 2016; Kravitz et al. 2013; DiCarlo et al. 2012). This is a result of processing that develops across a hierarchy of visual areas in the brain, building both selectivity and invariance at each stage.

At the early stage of the hierarchy, in primary visual cortex, selectivity and invariance are often thought of in terms of a dichotomy between simple and complex cell types (Hubel and Wiesel 1962; Skottun et al. 1991). Both cell types are typically selective for oriented features that are normally found in edges or gratings, which can be described as Gabor-like features containing interleaved stripes of light and dark regions (see filter in Fig. 1a). While simple cell responses are highly selective to a particular spatial alignment of these stripes within their receptive fields (RFs), complex cell responses are far more invariant to shifts in the alignment of the stripes across their receptive fields (Hubel and Wiesel 1962). This distinction has frequently been assessed by measuring each cell’s response to drifting sinusoidal gratings at its preferred orientation and spatial frequency, whereby simple cells show a modulated response to the drift, while complex cell responses are far less modulated (i.e. invariant) (Movshon et al. 1978a, 1978b; Dean and Tolhurst 1983). This dichotomy of cell types in primary visual cortex is supported by studies showing a bimodal distribution of the modulation ratio used to characterise responses to drifting gratings (Skottun et al. 1991).

**Figure 1.**
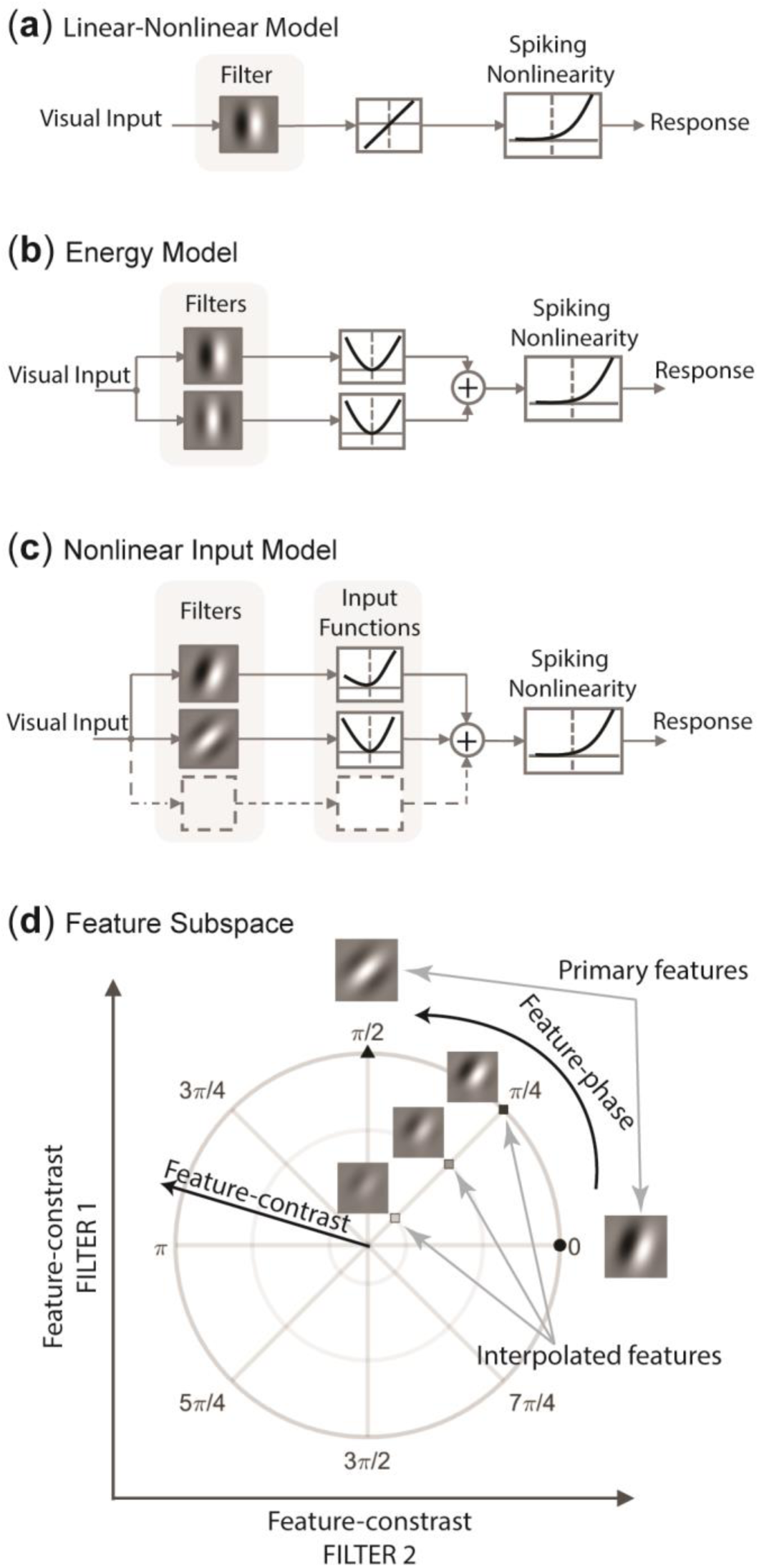
(**a**) Schematic diagram of the Linear-Nonlinear model used to describe simple cells. (**b**) Schematic diagram of the Energy Model used to describe complex cells. (**c**) Schematic diagram of the Nonlinear Input Model used for receptive field identification. (**d**) The feature subspace conceptualised as the linear subspace spanned by the primary stimulus features for a cell. Having described the subspace using polar coordinates, the angular direction, referred to as feature-phase, indicates different linear combinations (i.e. interpolations) of the two relevant features, and the radial direction indicates different feature-contrasts for each interpolated feature.

The dichotomy is further reflected in the standard models used to describe simple and complex cells. For simple cells a two-stage linear-nonlinear (LN) model is the standard (Fig. 1a), consisting of linear filtering with a Gabor-like filter spatially aligned to the cell’s receptive field, followed by a half-rectifying-like nonlinearity to describe the conversion to a spike rate. The linear stage captures the linear spatial summation properties emphasised in the original definition of simple cells (Hubel and Wiesel 1962). For complex cells the Energy Model depicted in Figure 1b is the standard, the first stage of which consists of two Gabor-like filters with identical orientation and spatial frequency, but separated in spatial phase by 90° (Adelson and Bergen 1985; Emerson et al. 1992). In the second stage, the filter outputs are squared and summed to give the response (equal to the “energy” in the orientation and spatial frequency bands). The LN and Energy Models respectively predict modulated (i.e. selective) and unmodulated (i.e. invariant) responses to drifting gratings.

Other studies have questioned the binary classification of neurons in primary visual cortex in terms of either simple or complex types. Some have argued that the cell population exhibits a continuum of modulation ratios in response to drifting gratings (Mechler and Ringach 2002; Hietanen et al. 2013; Cloherty and Ibbotson 2014; Meffin et al. 2015). Other studies directly estimated various filter-cascade style models designed as generalisations that encompass both the LN and Energy Models, by allowing variable numbers of filters in the model whose output could be pooled in some nonlinear fashion to produce a response. The models were estimated by applying statistical inference techniques, such as spike triggered covariance (Schwartz et al. 2006; Sharpee 2013) to large datasets of responses to stimuli such as white Gaussian noise or natural scenes. While these studies generally agree that these generalised models, when estimated directly from experimental data, exhibit a resemblance to the standard models, most have also noted important discrepancies (e.g. Rust et al. 2005; Chen et al. 2007; Fournier et al. 2014; for exceptions see Touryan et al. 2002, 2005). Some studies have suggested that the standard models need revision (Rust et al. 2005; Fournier et al. 2014; Carandini et al. 2005; Olshausen and Field 2005). For example, several studies have found that the estimated models of both simple and complex cells frequently contain more RF filters than exist in the corresponding standard models (Rust et al. 2005; Fournier et al. 2014). Furthermore, these RF filters can differ between themselves in orientation, spatial frequency or other characteristics, which is not expected in the standard models. This highly quantitative approach of direct model estimation reveals that both simple and complex cells include forms of nonlinear processing not accounted for by the standard models. The approach also reveals a greater diversity of processing in the cell population within the primary visual cortex than considered in standard models, consistent with a some earlier, more qualitative work that proposed more diverse classification schemes (e.g. Ikeda and Wright 1975; Henry 1977; Pollen et al. 1978).

A consequence of this diversity is that the models estimated in some of these studies can exhibit novel forms of invariance not captured by the standard models. These include invariance to perturbations in orientation and spatial frequency of Gabor-like features, in addition to the standard spatial phase invariance of complex cells (Chen et al. 2007). They can also possibly provide a more mechanistic description of how selectivity and invariance arise in the model. However, a description of these novel forms of invariance at the population level in primary visual cortex has not been previously reported. This has been hindered by a number of computational issues and constraints in estimating receptive field models for these cells. Our knowledge about the nonlinear feature selectivity and invariance of cells in primary visual cortex is primarily based on applying the spike-triggered covariance technique (Touryan et al. 2002, 2005; Rust et al. 2005; Chen et al. 2007). This method was one of the first to allow estimation of the RF filters in a multi-filter model, however, it suffers from some shortcomings and constraints.

First, as pointed out in several studies (Paninski 2003; Sharpee et al. 2004; Schwartz et al. 2006), the spike-triggered covariance technique is prone to biased or artefactual representations when used in conjunction with non-Gaussian stimuli. Later, several RF characterisation methods were put forward that allowed unbiased estimation of cortical RFs to arbitrarily distributed stimuli, amongst which are methods based on information theoretic measures (Sharpee et al. 2004; Kouh and Sharpee 2009; Fitzgerald et al. 2011a, 2011b; Rowekamp and Sharpee 2011; Rajan et al. 2013), minimising model prediction error (Rapela et al. 2010), and maximum likelihood estimation (Park and Pillow 2011; Kaardal et al. 2013; McFarland et al. 2013; Park et al. 2013). Second, the estimation of the nonlinearities applied in the second stage of the model following the initial filtering runs foul of the curse of dimensionality. That is, the number of parameters required to describe a completely general nonlinearity grows exponentially with the number of filters, making it impractical to estimate for more than two filters. Other methods have attempted to overcome the problem by making assumptions about the form of the nonlinearity employed by cortical neurons (Rust et al. 2005; Park et al. 2013; Rajan et al. 2013), which effectively reduces the dimensionality of the parameter space. These assumptions have varied in their degree of constraint and biological plausibility. Furthermore, almost all characterisation methods have represented the RF filters of cortical cells as a set of orthogonal filters in a vector space. This orthogonality constraint, which is there to facilitate computation of the RF filters, can lead to misleading interpretations. Kaardal et al. (2013) advocated that lifting this orthogonality constraint can be advantageous when inferring the mechanism underlying neural computation.

Amongst the approaches used for RF model estimation, the nonlinear input model (NIM) estimated using the maximum likelihood method provides a biologically plausible framework to describe single cell responses with minimal constraints on the form of nonlinearity (and filters), while using a description for which the number of parameters scales linearly with the number of filters (McFarland et al. 2013). Previous methods for estimating the NIM have fitted parameters sequentially for the linear and nonlinear stages of the model, a procedure that can lead to biased estimates (Rowekamp and Sharpee 2011). To avoid this problem, here we developed a procedure for the joint estimation of all parameters in the NIM via a maximum likelihood optimisation that can find the global maximum despite the presence of other local maxima.

Our aim is to objectively model the mechanisms underlying early cortical visual processing based on our improved method for estimating the NIM. Using recordings from a population of neurons in response to spatially white Gaussian noise, we reveal the full range of image features to which cells are sensitive through their RF filters and show that there is a range of nonlinear pooling mechanisms in primary visual cortex that increase the capacity for invariance to feature characteristics such as orientation, spatial frequency and spatial phase.

## Methods

### Preparation and surgery

Extracellular recordings were made from primary visual cortex in six anesthetized cats using methods described previously (Meffin et al. 2015). We recorded from Area 17, which is classed as primary visual cortex (Payne and Peters 2001). We refer to our recording site throughout the paper as primary visual cortex (V1). Experiments were conducted according to the National Health and Medical Research Council’s Australian Code of Practice for the Care and Use of Animals for Scientific Purposes. All experimental procedures were approved by the Animal Care Ethics Committee at the University of Melbourne (ethics ID 1413312).

Briefly, adult cats (2-6 kg) were induced with an intramuscular injection of ketamine hydrochloride (20 mg/kg i.m.) and xylazine (1 mg/kg), intubated, cannulated, and placed in a stereotaxic frame. Once intubated, oxygen and isoflurane were used to maintain deep anaesthesia during all surgical procedures. A craniotomy was performed to expose cortical areas 17 and 18. Anaesthesia was switched to gaseous halothane during data recording (0.5-0.7% during recordings) and the depth of anaesthesia was determined by monitoring a variety of standard indicators. To avoid eye movements during recordings, muscular blockade was induced and maintained with an intravenous infusion of Verucronium Bromide (i.e. Norcuron) through the infusion line at a rate of 0.1 mg·kg^-1^·h^-1^. Mechanical ventilation was utilized to maintain end-tidal CO_2_ between 3.5% and 4.5%. After an experiment the animal was humanely killed with an intravenous injection of an overdose of barbiturate (pentobarbital sodium, 150 mg/kg) while under anaesthesia. Animals were then perfused immediately through the left ventricle of the heart with 0.9% saline followed by 10% formol saline and the brain extracted.

### Visual stimuli and data recording

Visual stimuli were generated using a ViSaGe visual stimulus generator (Cambridge Research System, Cambridge, UK) on a calibrated, Gamma corrected LCD monitor (ASUS VG248QE, 1920×1080 pixels, refresh rate 60 Hz, 1 ms response time) at a viewing distance of 57 cm. White Gaussian noise stimuli comprising 90×90 pixels over 30° of the visual field were generated and employed to estimate the Neural Input Model (see ***Methods*** below). This allowed a 3-pixel resolution over 1° of the visual field, which provides a sampling rate of 3 cycles/degree. Given that having a cell with a spatial frequency preference of higher than 1.5 cycles/degree in cat V1 is very uncommon, the spatial resolution used minimised the risk of aliasing in estimating the visual receptive fields. The white Gaussian noise used in the stimuli had a mean value equal to the mid-luminance of the display monitor and a standard deviation chosen to result in a 10% saturation rate for individual pixels, i.e. the mean had a normalised intensity of 0.5, and 10% of pixels had a value of either 0 or 1 corresponding to the lowest and highest luminance of the monitor. Each noise frame was presented for 1/30 sec, followed by a blank screen of the mean luminance (intensity = 0.5), displayed for the same duration in blocks of 12,000. The blank period aimed to increase the overall response of the cell to the stimuli by increasing the temporal contrast.

Drifting sinusoidal gratings were also presented with 100% Michelson contrast, in a circular aperture and on a grey background of mean luminance. Various combinations of spatial and temporal frequencies were presented to determine the preferred directions of cells across the multi-electrode probe.

Extracellular recordings were made with single shank probes with Iridium electrodes (linear 32-electrode arrays, 6 mm length, 100 μm electrode site spacing; NeuroNexus), which were inserted vertically using a piezoelectric drive (Burleigh inchworm and 6000 controller, Burleigh instruments, Rochester, NY). Extracellular signals were acquired from 32 channels simultaneously using a CerePlex acquisition system and Central software (Blackrock Microsystems, Salt Lake City, Utah) sampled at 30 kHz and 16-Bit resolution on each channel. Filtering was performed by post-processing.

### Post-processing and spike sorting

Spike sorting of recordings was performed using KiloSort (Pachitariu et al. 2016), which is a system of spike sorting for dense electrode arrays. First, the raw data were band-pass filtered using a 3rd-order Butterworth filter, with lower and upper cut-off frequencies of 500 Hz and 14,250 Hz, respectively, in forward-backward mode to account for filter delay. Spikes were then detected using a dual-threshold algorithm to minimize the effect of small noise events. For a waveform embedded in the recorded time series to be detected as a spike, it had to satisfy two criteria: (1) every point (sample) had to exceed a weak threshold and (2) at least one point had to exceed a strong threshold. The weak and strong thresholds used were 3 and 4.5 times the standard deviations of the filtered signal, respectively. Using the geometry of the recording probe, an adjacency map was created that enabled detection of neighbouring points. This allowed clustering of spikes appearing simultaneously across multiple neighbouring channels as one unit. The spike clusters were manually curated for possible adjustment and verification using the graphical user interface *phy* (Rossant et al. 2016). Single units were identified during the manual curation process as units that manifested themselves as well-separated clusters in the feature space and exhibited a profound refractory period in their inter-spike interval histograms.

### Model estimation

#### Model Definition

We have adapted the NIM (McFarland et al. 2013) to improve the quality of fitting obtained to recordings from neurons in cat V1. The model diagram is depicted in Figure 1c, and describes the firing rate of the cell as a function of the input visual stimulus (McFarland et al. 2013),

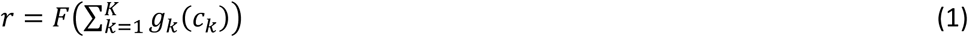

where *c*_*k*_ = **h**_*k*_ · **s** is the *feature-contrast* of the stimulus **s** with respect to the spatial filter **h**_*k*_, which is defined as their inner product. The model cell conceptually sums inputs from a number of parallel synaptic input streams, *K*, to give a generator potential 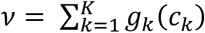. Each input is determined by an arbitrary function *g*_*k*_(·) (termed *input function*) of the feature-contrast *c*_*k*_ of filter **h**_*k*_, which captures processing performed by one or more presynaptic neurons. The number of input streams (i.e. RF filters) for each cell is determined using a statistical significance test described in the following subsection. The function *F*(·) indicates the overall spiking nonlinearity of the cell that converts the generator potential into firing rates and is described using a parametric representation as described in the following subsection. The response given by composition of the function *F*(·) with the summed input functions 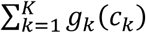 of the individual feature-contrasts, is the termed feature-contrast response function, which acts on the K-dimensional space of feature-contrasts (see Fig. 1d). The model assumes that the responses **R**_obs_ = {*R*^(1)^, …, *R*^(*T*)^} (integer spike counts) to the presented set of mutually independent stimuli *S* = {**s**^(1)^, …, **s**^(*T*)^} follow a homogenous Poisson distribution function

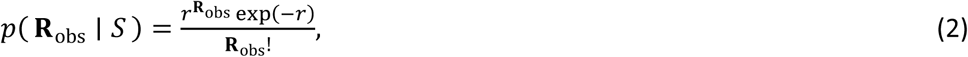

where *r* is the firing rate function described in Eq. 1.

#### Model Representation

Spatial RF filters were represented as

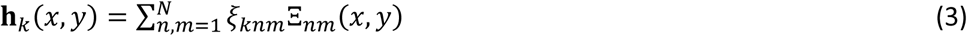

using a two-dimensional (2D) Fourier basis set of the form

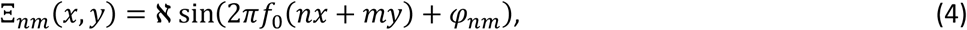

where *n* and *m* are integers from the set {–*N*, –*N* + 1, …, *N* – 1, *N*}, and *φ*_*nm*_ is a phase chosen to be either sine phase (*φ*_*nm*_ = 0 for (*m* > 0)) or (*m* = 0 & *n* > 0)) or cosine phase (*φ*_*nm*_ = *π*/2 for (*m* < 0)) or (*m* = 0 & *n* ≤ 0)). The conditions on *n* and *m* ensure there is no degeneracy in the basis. (*x, y*) are the horizontal and vertical pixel locations, *f*_0_ is the base spatial frequency and ℵ is a normalisation factor. We chose *N* = 5 to give 11 × 11 = 121 basis functions. This gave a representation of arbitrary spatial filters containing spatial frequencies up to *Nf*_0_, which was calibrated to approximately match the upper cut-off spatial frequency of the cell (= 10% of maximum amplitude, see *Feature Characterisation* below), and covering 2.3 octaves to the lowest sampled spatial frequency (other than 0). A square spatial region of interest was selected to enclose each cell’s receptive field filters spanning a single cycle of the lowest sampled frequency (i.e. of side 1/*f*_0_). This choice also resulted in the basis functions being orthonormal, which allowed straightforward computation of filter outputs from their coefficients, *ξ*_*knm*_, by converting stimuli, such as white Gaussian noise, into the Fourier basis representation in Eq. 4. This representation was found to give superior fits compared to pixelized representations because it resulted in a dimensionality reduction that reduced the number of parameters in most cases and eliminated high frequency noise from the fitting processes.

Input functions for each filter were represented as piecewise linear functions of the feature-contrast as 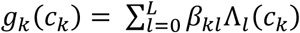 using a set of tent basis functions of the form (McFarland et al. 2013),

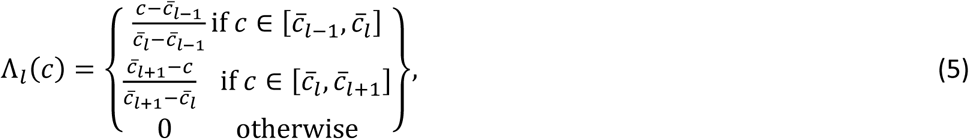

with *L* = 7 equally spaced intervals 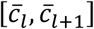 chosen to cover the 2.5% to 97.5% interval in the distribution of feature contrast, *c*_*k*_. In general, an arbitrary continuous function on a finite interval may be approximated as piecewise linear such that the mean-squared error reduces to zero as the number of intervals increases. *L* = 7 equally spaced intervals provided reasonable fidelity with just 7 parameters (*β*_*kl*_) per input function. To obtain the best possible fits to data, we did not place any constraints on the input functions (such as requiring them to be positive or negative to reflect excitatory or inhibitory inputs, respectively).

The spiking function that maps generator potential, *v*, to mean spike rate, *r*, was modelled as a log-exponential function 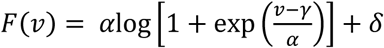. This function transitions from a constant spontaneous spike rate of *δ* ≥ 0, when the generator potential *v* is significantly below the threshold 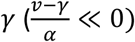 to a linearly increasing function with unit gain when the generator potential *v* is significantly above the threshold 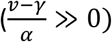. The transition occurs over a range of spike rates determined by *α* > 0 for which the function has a strictly convex shape.

#### Estimating global maxima

The estimation of the NIM parameters was achieved by maximising the log-likelihood of the model in Eq. 2, given the presented stimuli to the cell and their evoked responses:

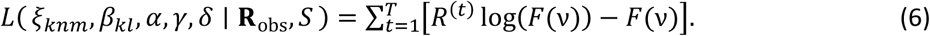

Previous applications of the NIM have used a sequential procedure to optimise the parameters (McFarland et al. 2013), typically in the following order: filters, input functions, spiking function-while holding the other parameters fixed at each stage (i.e. *ξ*_*knm*_ for **h**_*k*_, *β*_*kl*_ for *g*_*k*_ and (*α, γ, δ*) for *F*, respectively). We found that this sequential procedure tends to preserve the type of input functions that were chosen to initialise the model: i.e. if type-x functions are used initially, then the final input functions also tend to be of type-x and the filters are optimised to give the best match to input functions of this type, where type-x functions could be linear, threshold-linear, quadratic, etc. Similar problems with sequential optimization have been noted previously (Rowekamp and Sharpee 2011). We wish to avoid this arbitrary selection of the type of input function and associated filters, and instead find its optimal form over all model parameters simultaneously (see below).

We found that most cells had multiple local optima but observed that these optima all tended to be located in the same low dimensional subspace of RF filters of the cell (dimension ≤ 5). Due to the low dimensionality of this filter subspace, it was feasible to perform a systematic brute-force search over parameter values that covered this space. This search was only necessary for cells with multiple filters, as cells with just one filter have a unique optimal filter and feature contrast response function. An overview of the fitting procedure is as follows:

> Stage 1. Identifying the optimum number of RF filters and the corresponding subspace of RF filters. Other model hyperparameters, such as regularisation factors to prevent overfitting, were also optimised. These steps were done using a cross-validation procedure.
>
> Stage 2. Global optimisation using a brute-force search based on the subspace of RF filters found in Stage 1.

Conceptually, the different local optima correspond to filters (i.e. vectors) that point in a different set of directions in the same subspace (i.e. they form a different basis set for the same subspace). The corresponding input functions correspond approximately to different cross-sections through the feature-contrast response function in the direction of their filter - strictly to different cross-sections through the generator potential prior to application of the spiking nonlinearity. The two-stage optimisation procedure summarised above finds those filter directions in the subspace that allow a maximally separable generator potential *v* within the feature-contrast response function (i.e. the generator potential *v* can be written as sum of 1-dimensional (input) functions of distinct variables, i.e. feature-contrasts).

##### Stage 1: Identifying the subspace of RF filters

Optimising all model parameters simultaneously (for filters, input functions and spiking nonlinearity) from random or arbitrary initial values typically resulted in poor fits. It was therefore necessary to carry out Stage 1 using a sequential optimisation, using the following sequence: filters, input functions, spiking nonlinearity followed by filters again, holding other parameters fixed at each step. In the first step of this sequence, when optimising the filters, the parameters of the filters, *ξ*_*knm*_, and spiking nonlinearity, (*α, γ, δ*), were initialised randomly. For the input functions, it was necessary to choose a fixed parametric form. We used three different choices, to give three different model fits that were each carried through the entire two stage optimisation procedure. These were quadratic input functions (*g*(*x*) = *wx*^2^), threshold-linear input functions (*g*(*x*) = *w*[*x*]_+_) and a mixed model containing one linear input function (*g*(*x*) = *wx*) with the remainder quadratic (in these functions *w* determines the scale and sign of the input function, and was effectively fitted during the first step as the norm of the filter, ‖**h**_*k*_‖). We found that the final optima estimated for different models were highly similar provided the full two stage global optimisation was used.

In the second step of the sequence, the input functions were initialised as piecewise-linear approximations to the input functions from the first step (i.e. quadratic, threshold-linear or mixed) prior to optimization of their parameters, *β*_*kl*_. For all other parameters and subsequent steps in the sequence, optimisation began from the parameter values obtained at the end of the previous step.

As the complexity (number of parameters) of a model increases, it becomes more susceptible to overfitting the data. Hence, it is crucial in model estimation to take necessary measures to prevent a model from overfitting. We employed cross-validation to avoid overfitting and determine the hyperparameters of the model, including the number of filters.

The number of filters for each cell was systematically varied while the statistical significance of each filter was evaluated by bootstrapping (Supplementary Fig. 1). For this, we divided the data into a training set, which comprised four-fifths of the data, and a test set, which comprised the other one-fifth of the data. For each specified number of filters, we used the training set to estimate the filters, and then assessed the performance of the model by computing its log-likelihood using resampling from the test set (this was repeated 500 times). Thus, for each number of filters, we found a distribution for the log-likelihood computed on the test set. The inclusion of a new filter was counted as significant if it significantly improved the log-likelihood of the model on the test set (Z-score > 2).

In Stage 2, regularisation was used as another measure to prevent the input functions from overfitting to data. For this, we enforced regularisation on the input functions by penalizing their second-order derivatives. The weight by which the regularisation was added to the log-likelihood of the model (i.e. objective function) was determined as a final step in Stage 1 using five-fold cross-validation, in the same process as described above.

The end-result of Stage 1 was three different NIMs with filters, input functions and spiking nonlinearities optimised sequentially. Although the different NIMs were initialised with different input functions (quadratic, threshold-linear and mixed linear/quadratic) and had different optimal RF filters and input functions, the subspaces spanned by the optimal RF filters were substantially similar.

##### Stage 2: Brute force global optimisation

The filter subspace identified in Stage 1 was used to initialise optimisations in Stage 2 over all parameters. This form of initialisation allowed a local gradient ascent algorithm to find “good” local optima with high log-likelihood compared to a random initialisation. For cells with multiple filters, the filters initialised as different linear combinations of the normalised filters, 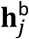, identified in Stage 1 to cover the subspace approximately uniformly: 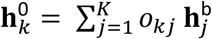. Different initialisations could lead to different local optima, allowing the global optimum over all initialisations to be selected. Generally, we found from one up to several local optima, with many initialisations converging to the same optima. The *o*_*kj*_ are parameters sampled from the upper hyper-sphere of dimension *K* – 1, so as to preserve the unit norm of the initialised filters 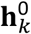 given the normalised basis filters 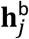 for the subspace; sampling only from the upper hypersphere avoids degenerate initialisations that are simply opposite in sign to previous choices. For example, models with two filters are described by polar coordinates *o*_*k*1_ = cos *θ*_*k*_, *o*_*k*2_ = sin *θ*_*k*_ with *θ*_*k*_ ∈ *mπ*/*M*│*m* = 0, …, *M* – 1 & *θ*_2_ > *θ*_1_ (to avoid repeats or degeneracy). For higher dimensions, we used *o*_*k*1_ = 1 and *o*_*kj*_ ∈ {–1, +1}, *j* ≠ 1 to ensure sampling from the upper hyper-sphere in the *K*-dimensional feature subspace (filters were then normalised). For each set of initial filters, we ensured that they were independent by excluding the cases where the transformation was singular. For models with three filters this initialisation yielded 16 different combinations of filter directions. For models with more than three filters, the number of all initial sets of filters grows combinatorially, which makes it computationally infeasible to probe them all. To deal with this, we randomly chose 30 initial sets for models with four filters, and 40 initial sets for models with five filters. All initial filters were normalised, and for each initialised set of filters a sequential optimisation over input functions *g*_*k*_(·) and the spike function *F*(·) was performed to initialise these functions to values close to their (simultaneous) optimum given the initial filters. Then a simultaneous optimisation was performed using the parameters *ξ*_*knm*_, *β*_*kl*_ and (α,γ,δ) for the filters, input functions and spike function, respectively. Note this includes optimisation over all filter parameters, and not just those that restrict the model to the subspace identified in Stage 1.

Also note that in cases where models included only one spatial filter, the input function and the spike function can be composed together and be described by a single function. Therefore, to avoid any degeneracy for models with one spatial filter, the spike function parameters (α, γ) were held fixed at (1,0), and the optimisation was performed over the rest of the parameters.

#### Avoiding model degeneracy

The form of the NIM with a generator potential given as the sum of input feature-contrast functions is intended to approximate synaptic integration as a linear process given nonlinear inputs from presynaptic neurons with different feature sensitivities (McFarland et al. 2013). The form described here, with entirely arbitrary input functions (i.e. not restricted in sign to be either excitatory or inhibitory), has the advantage that its representation is provably non-degenerate, meaning that there are no alternate parameter choices that can describe a functionally identical model. This is a consequence of the generator potential being an additively separable function of feature-contrast, and is true provided that four conditions are met for all values of *k*: i.e. (1) ‖**h**_*k*_‖ = 1, (2) *g*_*k*_(0) = 0, (3) *g*_*k*_(*c*_max_) > *g*_*k*_(*c*_min_), (4) 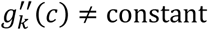 constant, where *c*_max_ and *c*_min_ are the maximum and minimum feature-contrast in the stimulus set, respectively. The first three conditions are arbitrary choices we implemented in the code to put the model in a standard form that would otherwise lead to degeneracy, while the final condition did not arise in our data set, but precludes models with strictly linear or quadratic input functions. This latter situation admits a continuous family of linear transformations on the filters, leaving the overall model invariant. Note also that the choice of a single parameter, *α*, in two places in the spike function (multiplying the logarithm as well as in the denominator of the argument of the exponential), avoids a degeneracy with the overall scale of the input function through the parameters *β*_*kl*_.

### Model analysis for feature selectivity and invariance

For each neuron fitted with the NIM, the model was analysed to describe the feature selectivity and invariance of the cell. This involved first defining the subspace of features to which the cell was sensitive, characterising and analysing those features, characterising the nonlinear response to those features, and finally quantifying the degree of response selectivity or invariance to the characteristics of those features.

#### Feature subspace and the interpolated features

The diagram in Figure 1c represents the RF model of a cell with multiple spatial filters. This cell is theoretically sensitive to any feature (i.e. spatial structure) in an image that is a linear weighted sum of the RF filters: these features will be referred to as the interpolated features as opposed to the features corresponding to the RF filters, which will be referred to as primary features. Note that here we are conceptually distinguishing features from spatial filters: the former are visual inputs embedded in an image, while the latter are mathematical operations that process images to quantify how much of the corresponding feature is present. Thus, primary features and their corresponding filters are conceptually different, despite having the same spatial structure. We define the subspace of features to which the cell is sensitive (i.e. its feature subspace) as that spanned by the primary features of the cell: it consists of the full set of interpolated features. Figure 1d shows how we present the feature subspace in polar coordinates. Different linear combinations of the primary features correspond to distinct points in the feature subspace. In the angular direction, referred to as *feature-phase*, the characteristics of the spatial form of the interpolated features can change. In the radial direction, the contrast of interpolated features vary, but not their spatial structure. Thus, to each feature-phase there corresponds a unique interpolated feature varying only in feature-contrast. The full set of interpolated features in the feature subspace, independent of feature-contrast, is referred to as the cell’s feature spectrum (e.g. the set of interpolated features on the unit circle).

#### Feature-contrast

The model considered in this study consists of a linear filtering stage in which a spatial filter **h** is applied on the stimulus **s**. The application of the filter to the stimulus is achieved by calculating the inner product **h** · **s** in a vector-wise manner, in which the output indicates the similarity between the visual stimulus and the spatial structure of the filter. This spatial filtering projects the stimulus onto the corresponding feature. The stimulus can be written as **s** = **s**_**1**_ + **s**_**2**_, where **s**_1_ is the component parallel to the filter and **s**_2_ is the component perpendicular to the filter **h**, i.e. **s**_1_ ∈ ℝ^n^: **s**_1_ ∥ **h** and **s**_2_ ∈ ℝ^n^: **s**_2_ ⊥ **h**. Therefore, **s**_1_ = *ρ***h** where *ρ* ∈ ℝ, and consequently **h** · **s** = *ρ* since the filter is normalised, i.e. ‖**h**‖_**2**_ = **1**. Assuming that the mean value of the feature is negligible, the RMS contrast of **s**_1_ would equal its norm *ρ*. This means that the output of the linear spatial filtering can be interpreted as the contrast of the feature corresponding to **h** embedded in the stimulus **s**, hence termed feature-contrast. Moreover, since **h** · **s** = ‖**s**_1_‖_**2**_ the contours of equal feature-contrasts in the feature subspace are concentric circles centred at the origin (i.e. zero feature-contrast). Here, for presentation purposes and to provide a more illustrative representation of the visibility of the embedded feature, we described the feature-contrast in terms of *Michelson* contrast.

#### Feature characterisation

We quantitatively characterised the features (and filters) associated with a cell model in terms of peak orientation, peak spatial frequency and relative spatial phase. These characterisations were performed non-parametrically, using the representation of the features in the Fourier domain (see Supplementary Fig. 2a, 2b). By representing the Fourier amplitude spectrum in polar coordinates, the angular and radial coordinates represent the orientation and spatial frequency contents of a feature, respectively. The peak orientation and peak spatial frequency of a feature correspond to the angular and radial coordinates of the feature’s peak amplitude spectrum, respectively. The spatial phase is defined as the phase difference between two features, simply calculated by taking the circular average of the Fourier phase over the spectral domain where amplitude exceeds half of its maximum (Supplementary Fig. 2d). For the interpolated features of a cell, the spatial phase is measured relative to the phase of the cell’s first primary feature.

**Figure 2.**
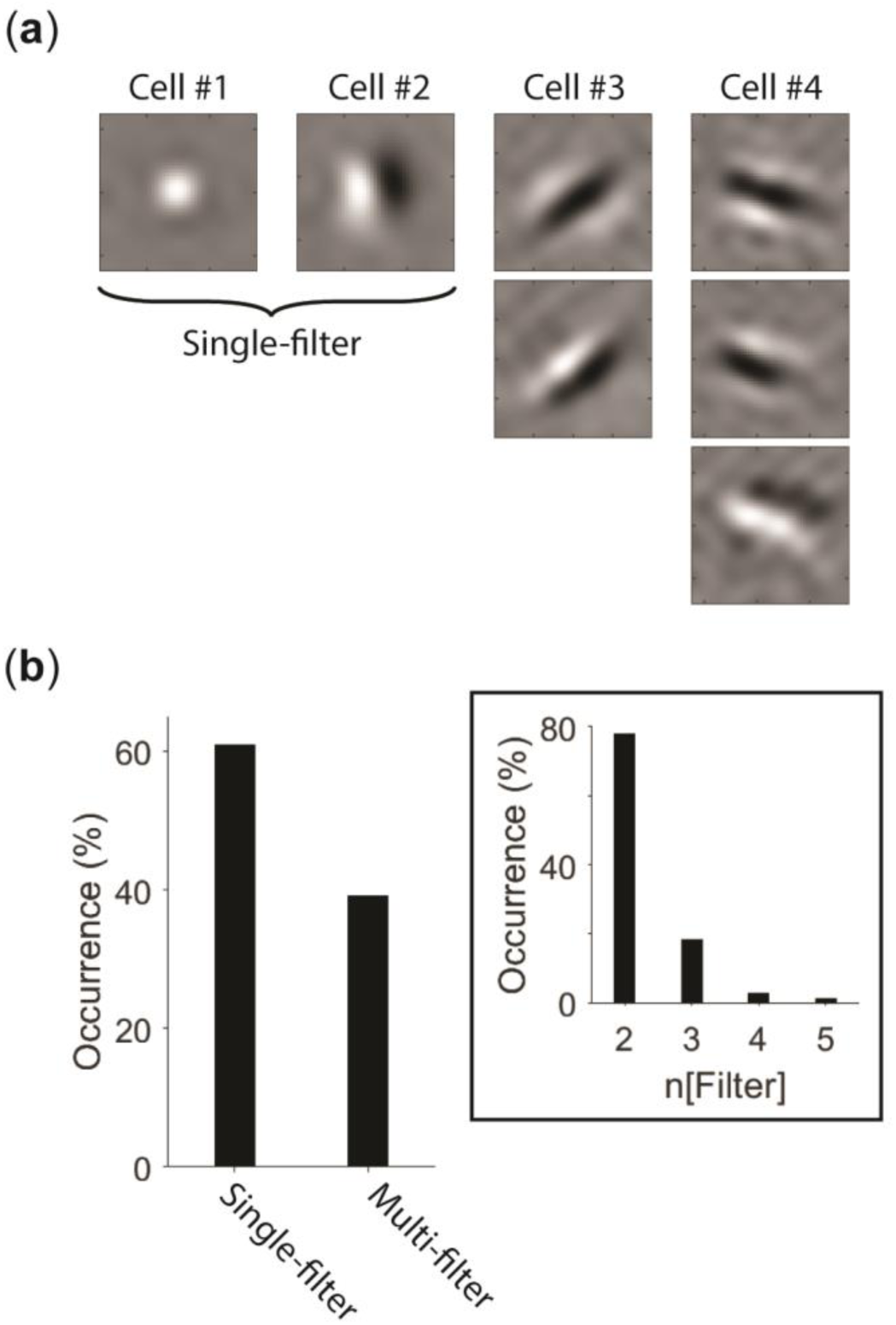
Characterisation of spatial feature selectivity of V1 cells. (**a**) Four example cells that pool across different numbers of spatial filters. (**b**) The bar graph presents the distribution of single-versus multi-filter cells in our population V1 data. The inset shows the distribution of identified numbers of filters for the multi-filter cells.

**Figure 3.**
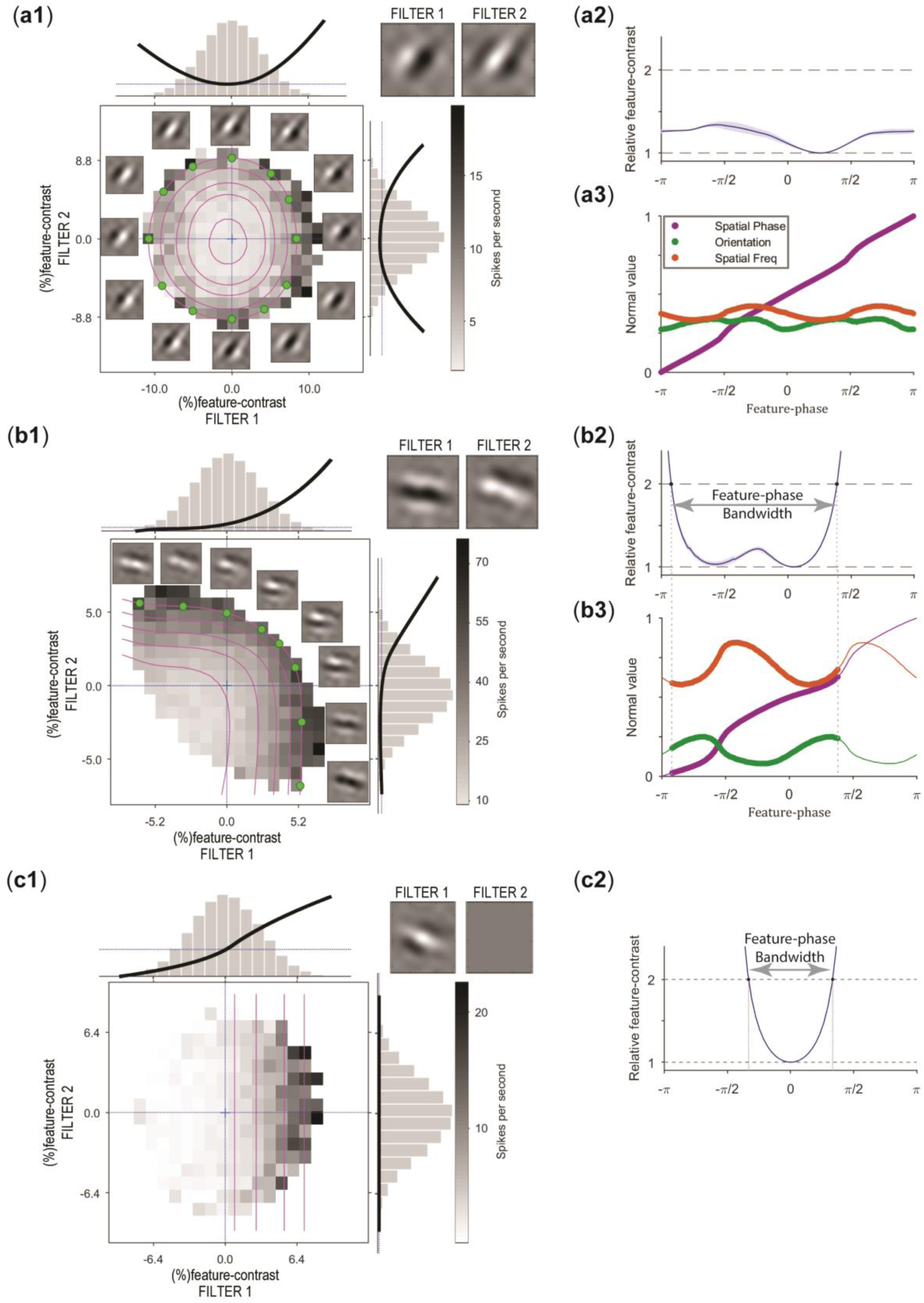
Example model fits to V1 cells. (**a1**) An example model fit to a cortical neuron for which we found spatial phase invariance (modulation ratio to drifting gratings 0.37). Grids indicate 2° of visual field. (Top right) The two identified spatial RF filters for this cell. The nonlinearities for its two filters are shown as black curves superimposed on grey bar-graphs, the latter showing the distributions of the WGN stimuli. The grey pixels show the empirically calculated mean responses of the cell across the WGN stimuli in feature subspace (conventions as in Fig. 1c). (Note that the white region surrounding the circular region of grey pixels corresponds to parts of the feature space not sampled by the WGN stimuli.) Each point in the 2D polar plot is associated with an interpolated feature consisting of a particular linear combination (i.e. feature-phase) of features specified by Filter 1 and 2 that corresponds to that pair of feature-contrasts (see **Methods**). The superimposed magenta iso-response contours show equal spike rates as functions of the feature-phase and feature-contrast (in terms of Michelson contrast). Some interpolated features associated with the locations of the green points are presented around the plot. (**a2**) The relative feature-contrast of the interpolated features along iso-response contours plotted as a function of feature-phase, averaged across different normalised spike rates. Shaded area = 1 standard deviation. (**a3**) Characteristics of the interpolated features at different feature-phases, sampled from an iso-response contour in the feature spectrum of the cell. The variation in each feature characteristic is normalised by its natural range of variation (90° for orientation; 0.6 cycles per degree for spatial frequency; 360° for spatial phase). (**b1**) Example model fit for a cell with two not even-symmetric input functions (modulation ratio 1.11). (**b2**) As a result of the openness of the iso-response contours for this cell, the relative feature-contrast varies significantly. An upper bound of 2 was set on relative feature-contrast to define feature-phase bandwidth (shown by the line with arrows). (**b3**) Characteristics of the interpolated features sampled from an iso-response contour in **b1**. The thick lines correspond to the interpolated features within the feature-phase bandwidth. (**c1**) An example model fit to a cell with one spatial filter (Filter 1). A dummy second filter (Filter 2) is considered to obtain a 2D representation of feature subspace for visualisation purposes. The linearity of the iso-response contours shows the dependency on only the feature-contrast of Filter 1. (**c2**) Due to the linearity of the iso-response contour plots, the relative feature-contrast of the interpolated features have the narrowest feature-phase bandwidth.

#### Feature classification

We classified spatial features into three types: Gabor-like, blob-like, and unclassified (see Fig. 4 for examples of these feature types). Qualitatively, Gabor- and blob-like features are distinguished by their spatial structures, i.e. showing significant alignment toward a particular orientation or lack thereof, respectively. To determine the type of feature quantitatively, we first calculated its representation in the Fourier domain, as described above. Then, we found all the significant maxima of the feature’s amplitude spectrum defined as those maxima exceeding half the global maximum amplitude, and which were separated from the global maximum by a dip in the amplitude spectra lower than half the global maximum. Ideally, an oriented Gabor-like feature, like the one shown in Supplementary Fig. 2a, has two significant maxima in its amplitude spectrum confined to localised regions at equal but opposite spatial frequencies (the two patches in red in Supplementary Fig. 2b). Our measure for classifying an oriented Gabor-like feature is the orientation bandwidth of the feature, defined as the full width at the half maximum (FWHM) of the amplitude spectrum along the circle passing through the peak amplitude spectrum (Supplementary Fig. 2c). As the orientation bandwidth of a Gabor-like feature increases, it becomes less oriented; eventually it reaches a transition point that it is no longer regarded as selective for orientation and transforms into a blob-like feature. An ideal blob-like feature has an orientation bandwidth encompassing the full circle of 360° in Fourier space. Here, we chose an orientation bandwidth of 120° as the threshold between a Gabor-like and a blob-like feature. Therefore, a feature is classified as Gabor-like if (a) it has two significant maxima in its amplitude spectrum, and (b) it has an orientation bandwidth <120°. A feature is classified as blob-like if it has an orientation bandwidth ≥120°. We also found unusual interpolated features that had more than two significant local maxima in their amplitude spectrum, which corresponded to multiple peak orientations or peak spatial frequencies, indicating they were not unique. The spatial structure of such features often resembled a three- or four-leaf clover pattern. The feature presented in Figure 4c is one such example: it shows a preference for two different (approximately perpendicular) orientations. These features do not fall into any of the Gabor-like or blob-like classes and are, therefore, labelled unclassified. As most cells had more than 90% Gabor-like features (Fig. 4a), only this type of feature was included in the analysis of feature selectivity and invariance. This allowed us to analyse selectivity and invariance in terms of the feature characteristics of peak orientation, peak spatial frequency and spatial phase which are well-defined for Gabor-like features.

**Figure 4.**
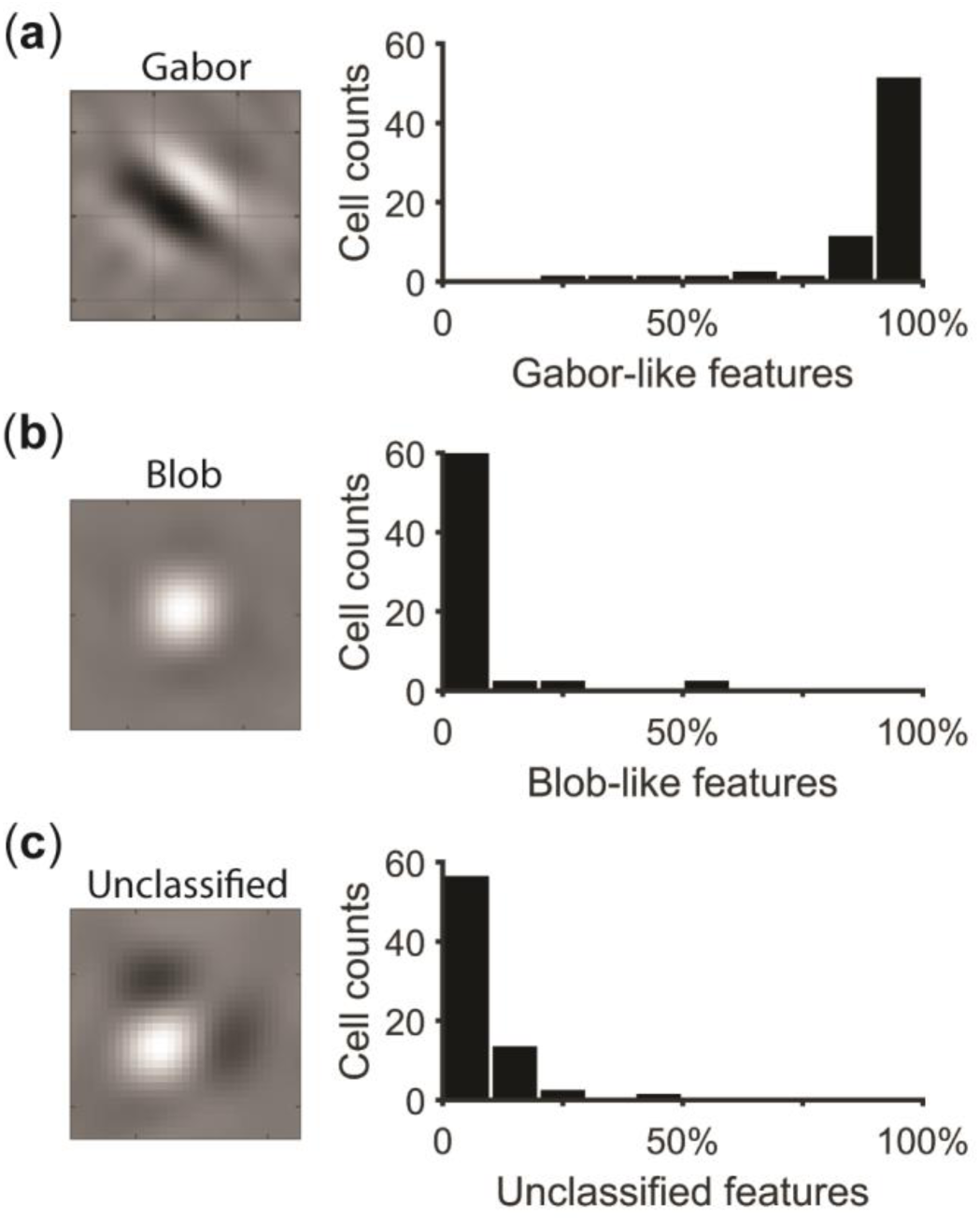
Distribution of different types of interpolated features within the feature spectra of the cells in our population of V1 cells. The features were classified into three types: (**a**) Gabor, (**b**) Blob, and (**c**) Unclassified. The spatial structures on the left show example features for each class. The bar graphs indicate the number of cells having a given fraction of feature type in their feature spectrum. Most cells have more than 90% Gabor-like features

**Figure 4.**
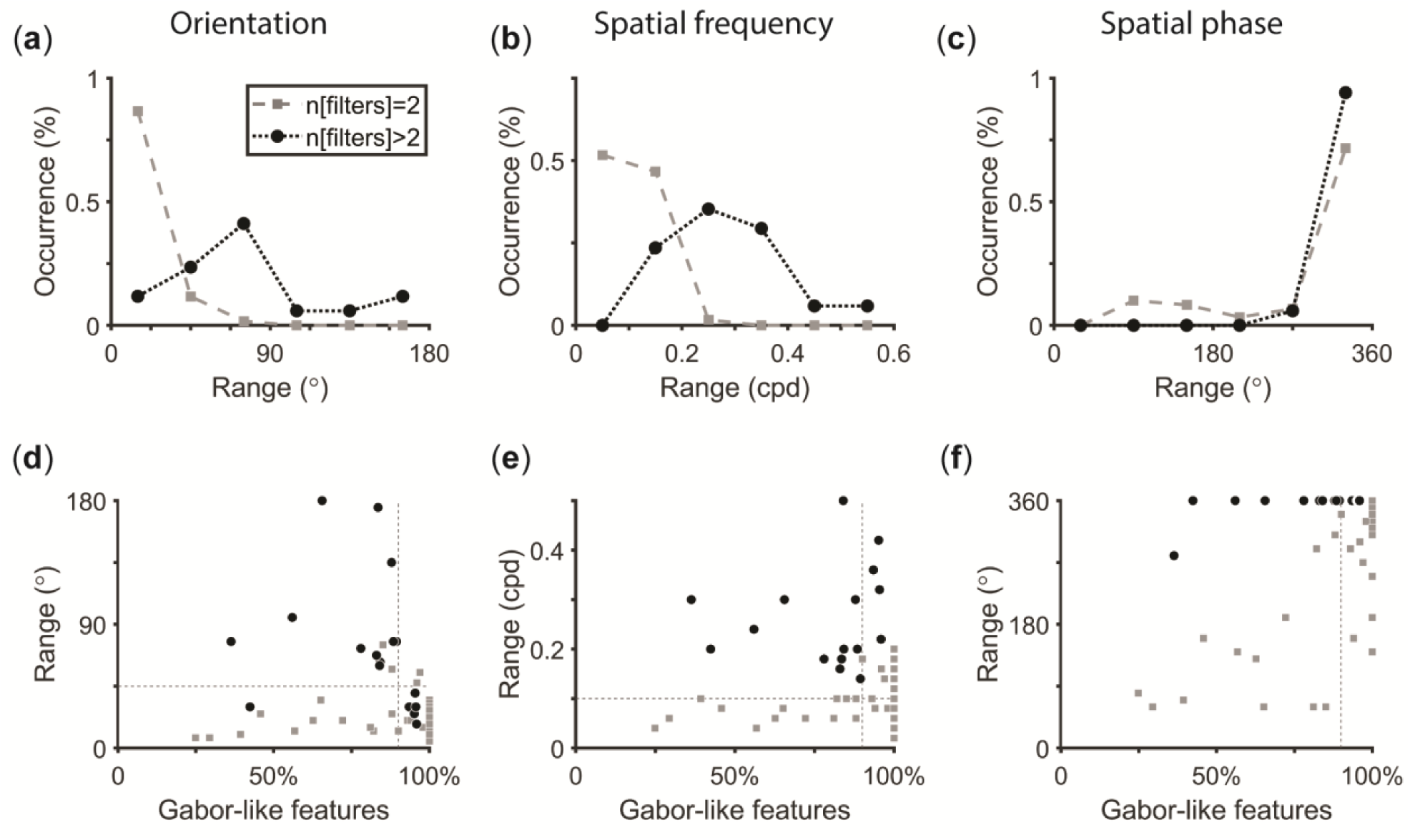
Feature spectrum characteristics of V1 cells. The top row plots the distributions of the spectral ranges in (**a**) orientation, (**b**) spatial frequency, and (**c**) spatial phase, spanned across the cells’ feature spectra. The distributions are split for cells pooling two features (dashed light lines, square symbols) and cells pooling three or more features (dotted dark lines, circle symbols). The bottom row shows scatter plots describing the spectral ranges for (**d**) orientation, (**e**) spatial frequency, and (**f**) spatial phase versus the fraction of Gabor-like interpolated features within their spectra. The vertical dashed lines indicate 90% Gabor-like features. The horizontal grey dashed lines in **d** and **e** indicate 45° and 0.1cpd, respectively.

#### Calculating the spectral range of feature characteristics

To quantify the *spectral range* of variation of each of these three feature characteristics across the feature spectrum, we first discretised the cell’s feature spectrum in terms of a set, Ψ, of feature-phases, with each phase uniquely associated with an interpolated feature. For cells with 2 filters we used a uniform sampling of feature-phase *ψ* ∈ [0,2*π*] with 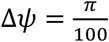. Using this discretisation technique, the set of feature-phases is described as Ψ = {*ψ*_1_, *ψ*_2_, …, *ψ*_*K*_}, where *ψ*_*k*_ = (*k* – 1)Δ*ψ, k* = 1, …, 200. For cells with 3 filters we used spherical polar coordinates, (*r*, θ, φ), with the latter two angular coordinates representing a 2-dimensional “feature-phase” *ψ* = (θ, φ) (and the first representing feature-contrast). In general, for cells with *N* > 2 filters we used the angular hyper-spherical coordinates for *ψ* = (θ, φ_1_, …, φ_*N*–2_), with generalised polar angles φ_*n*_ ∈ [0, *π*] and generalised azimuthal angle θ ∈ [0,2*π*] (so that in cartesian coordinates: 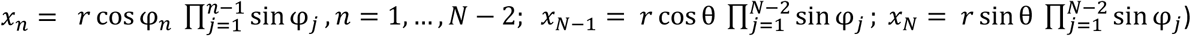. These angles were discretised with sampling steps dependent on the dimensionality: φ_*n,k*_ = (*k* – 1)Δφ, *k* = 1, …, *K* and θ_*n,k*_ = (*k* – 1)Δθ, *k* = 1, …, 2*K*, where 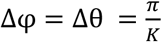 and the number of samples *K* = 50 for *N* = 3, and *K* = 20 for *N* = 4, 5.

For each discrete value, *k*, of feature-phase in the set Ψ, we calculated peak orientation, peak spatial frequency, and spatial phase of the corresponding interpolated feature. The spectral range of each feature characteristic was calculated as a sum over the characteristic values covered by the discrete set of feature-phases, Ψ. To do this, for each characteristic, *ϑ* ∈ {ori., freq., phase}, we computed its empirical histogram by binning the natural range of that characteristic as 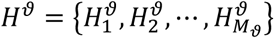, where 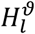 denotes the number of interpolated features whose *ϑ* characteristics fall into the *l*^*th*^ bin. *M*_*ϑ*_ is the total number of bins, and is determined as 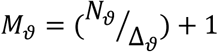, where *N*_*ϑ*_ indicates the maximal natural range and Δ_*ϑ*_ is the bin-size. The range of variation (*R*) for each feature characteristic is then calculated as the total width of bins containing at least one feature:

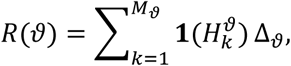

where

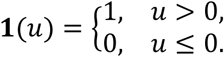

The bin sizes were chosen as Δ_ori_= 2.5°, Δ_freq_= 0.02 cpd and Δ_phase_= 5°. Also, the maximal natural ranges of variation that we considered were *N*_ori_ = 180°, *N*_freq_ = 0.8 cpd and *N*_phase_ = 360°.

#### Characterising input functions

The input functions were characterised using a symmetry index around the origin in feature-contrast. In general, a function *g* is said to be even-symmetric if *g*(–*x*) = *g*(*x*) for all *x* belonging to its domain, whereas an odd-symmetric function exhibits the property of *g*(–*x*) = –*g*(*x*). We calculated the symmetry index for each curve as

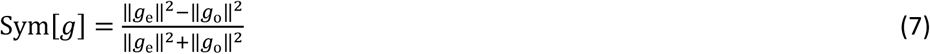

where *g*_e_ and *g*_o_ are the even and odd components of the function *g*, respectively, defined as

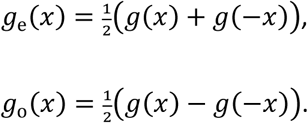

The operator ‖·‖ denotes the function norm in Hilbert space, defined as

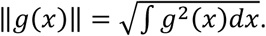

The symmetry index defined in Eq. 7 varies from −1, for an odd-symmetric function, to 1, for an even-symmetric function. A threshold-linear function results in a symmetric index of 0.

#### Calculating feature-phase bandwidth

To characterise the part of a cell’s feature spectrum over which it responds equally, we developed a measure referred to as the feature-phase bandwidth. It quantifies the portion of the cell’s feature spectrum effectively sampled by the feature-contrast response function (see Results for further intuition). To calculate this, we considered iso-response contours on a cell’s feature subspace, each giving a set of interpolated features for which responses were equal. We then calculated the feature-contrast and feature-phase of each interpolated feature, which gave the feature-contrast required to drive the cell at a given iso-response level as a function of feature-phase (for response levels ranging from 10% to 90% maximum response with 5% steps). To normalise these curves across different response levels, we divided by the minimal feature-contrast across all feature-phases on that contour, to obtain the *relative feature-contrast* as a function of feature-phase. The feature-phase bandwidth of the cell was defined as the range of feature-phases for which the relative feature-contrast, averaged across response levels, was less than a factor of 2. In other words, the feature-phase bandwidth gives the range of feature-phases with interpolated features capable of driving the cell at a fixed response level and with less than twice the contrast required by the feature to which the cell was most sensitive.

In practice, we use the discretisation of feature-phase described above (§ *Calculating the spectral range of feature characteristics*). For cells with two filters we used the set 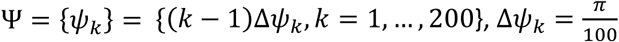. We calculated the feature-phase bandwidth as

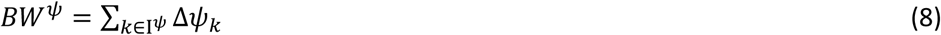

where the set 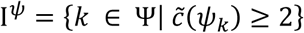 is the set of feature-phases indices for which the mean relative feature-contrast, 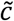, defined above, is less than 2. For cells with *N* > 2 filters, we used corresponding discrete sets, Ψ, of feature-phase based on the spherical or hyperspherical representation defined above (§ *Calculating the spectral range of feature characteristics*). For these cells, feature-phase bandwidth was generalised in terms of area, volume etc. on a unit hyperspherical surface centred on the origin, instead of a circle of 2*π* radians. This gives a discrete element of surface area on the hypersphere, 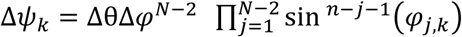, at each discrete feature-phase point *ψ*_*k*_ = (θ_*k*_, φ_1,*k*_, …, φ_*N*–2,*k*_), as defined above (§ *Calculating the spectral range of feature characteristics*). As the length/area/volume etc. of a unit hypersphere changes with dimension, we normalised the feature-phase bandwidth by this length/area/volume etc. to allow comparison across dimensions.

#### Calculating the tuning breadth of feature characteristics

The overall selectivity or invariance to each feature characteristic is a consequence of two factors: (1) the spectral range of that characteristic in the feature spectrum of the neuron, and (2) the feature-phase bandwidth that is determined by the pooling mechanism of the neuron in the form of the feature-contrast response function. We combined these two factors into a single measure quantifying the continuum between selectivity and invariance of response to a feature characteristic, which we refer to as the characteristic’s *tuning breadth*. This was done by considering the range of variation in the characteristics of interpolated features in that part of the cell’s feature spectrum covered by the cell’s feature-phase bandwidth defined in Eq. 8. This was done by calculating the range of variation in the characteristics of interpolated features with relative feature-contrast less than 2 (based on the definition of the feature-phase bandwidth).

#### Linear and quadratic model comparison

To further understand the role of the feature-phase bandwidth in determining the tuning breadth of the characteristic, (i.e. the second factor in the above section) we compared the tuning breadths obtained through the fitted NIM to two hypothetical alternative models that had either maximal or minimal feature-phase bandwidths, but applied to the same feature spectrum of the NIM of the neuron. The maximal model corresponds to a feature-contrast function with fully enclosed iso-response contours having a feature-phase bandwidth of 2π radians (e.g. circular iso-response contours formed from quadratic input functions). The minimal model corresponds to a feature-contrast response function that depends on only one feature dimension, and represents linear spatial processing and pooling, for the single feature that drove the cell at fixed rate using minimal feature-contrast. The model is the *r* = *F*(**h**_min_ · **s**), where **h**_min_ is the filter corresponding to the minimal feature and we have collapsed the input function into the spiking non-linearity to avoid degeneracy. This model has planar iso-response contours orthogonal to **h**_min_ and has a feature-phase bandwidth of 2π/3 radians.

### Calculating each unit’s modulation index

We measured the responses of single units to sinusoidal drifting gratings. The parameters of the gratings were chosen to correspond to the preferred tuning of the majority of units recorded in a given track. To calculate a modulation index, we performed Fourier analysis using the FFT function in Matlab^©^ on the spike train. From the Fourier spectrum we extracted the amplitude of the fundamental frequency (F1) and divided it by the mean response (F0), after subtracting the mean spontaneous activity from both measures (exactly the same method was used in Crowder et al. 2007). The F1/F0 ratio is the modulation index for that cell. The classical definition of a complex cell is that it has an F1/F0 ratio <1, while a simple cell has an F1/F0 ratio >1 (Skottun et al. 1991).

## RESULTS

### Part 1. Spatial receptive field analysis

We applied the NIM to recordings from neurons in cat V1 in response to spatially white Gaussian noise. The definition of terms is summarised in Table 1. The estimated RF filters for each cell (Fig. 2a) act on image stimuli to determine the primary image features to which the cell shows sensitivity. The primary image features have an identical spatial structure to the filters. Figure 2a presents example RF filters from four recorded V1 cells. Dark regions show where the cells responded to image luminance darker than the mean (OFF responses) and white regions show where the cells responded to luminance brighter than the mean (ON). In all cases, the filters were spatially localised. Cell#1 was selective for blob-like features of the appropriate brightness polarity that were presented within the filter’s spatial receptive field. The other example cells were selective for elongated Gabor-like features with alternate ON and OFF polarities. Cell#2 had a single filter, Cell#3 had two spatial filters with the same orientation but phase shifted by 90°, and Cell#4 had three filters. We recorded from 120 units with single filters (blob or elongated) and 77 units with multiple filters (Fig. 2b). For multi-filter cells, a majority (78%) had two filters, 18% had three filters and the remaining 4% had 4-5 filters (Fig. 2b; inset).

**Table 1.**
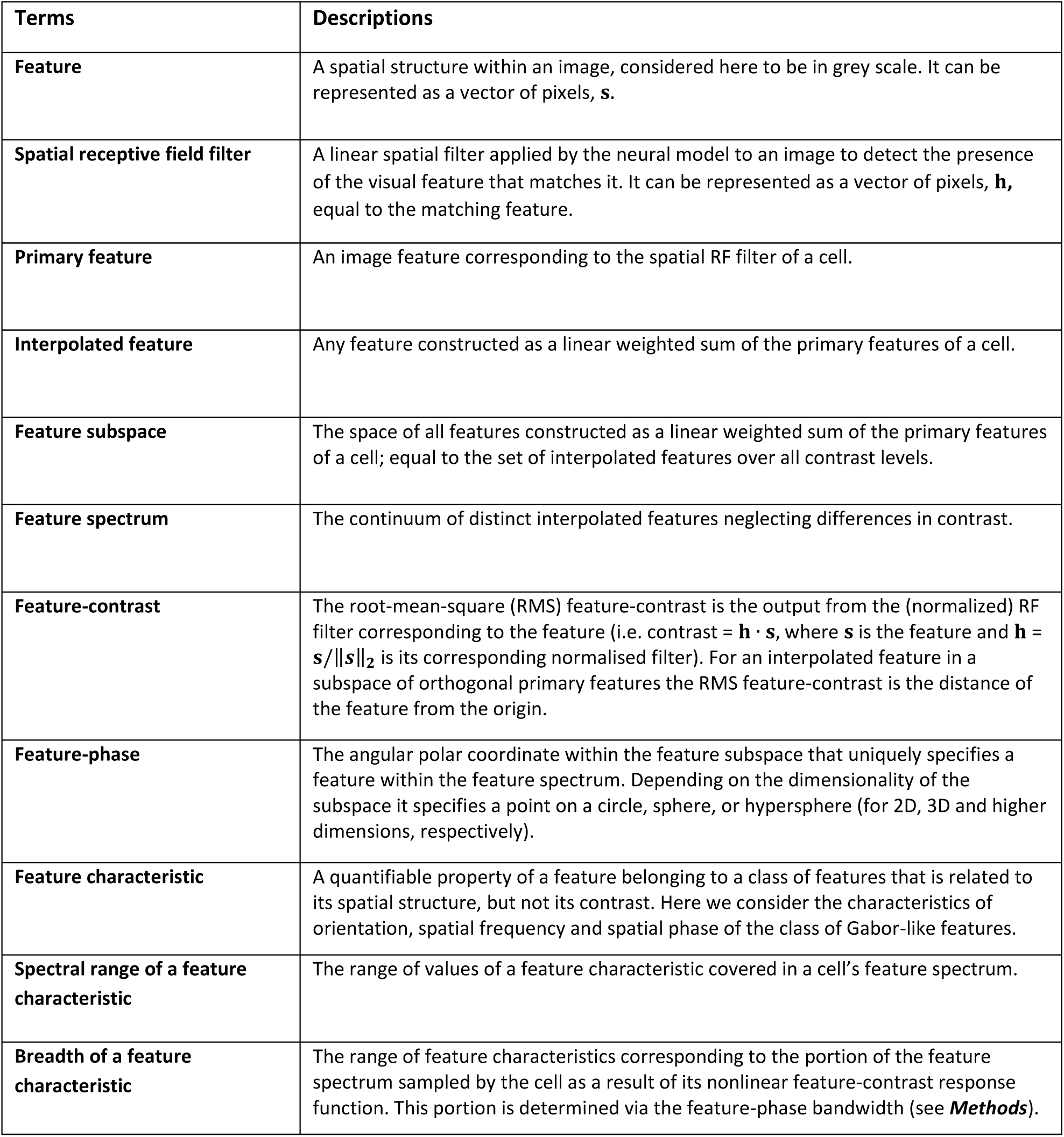
Definition of the terms used throughout this study.

### Part 2. Example cells

A cell’s selectivity or invariance to image features depends on the spatial structure of the filters, the relationship between filters and the form of the input functions (Fig. 1b). This section will illustrate the influence of these factors for three example cells that have at least 2 filters.

Figure 3 presents the three example cortical cells, with the estimated RF filters (corresponding to the primary features) shown in the top-right of each cell’s feature subspace. The feature subspace is the space that includes all possible linear weighted sums of the cell’s primary features: these summed features are referred to as the interpolated features (Fig. 1d). In Figure 3a1, we plot the mean spike rates (as shades of grey) in response to every interpolated feature in the feature subspace. The axes of this 2-dimensional subspace are represented by the feature-contrast of each primary feature embedded in the white Gaussian noise, and is equal to the output for the corresponding (normalised) filter (refer also to Fig. 1d). The curves in magenta indicate iso-response contours for several response levels. Adjacent to each axis of the subspace plot, we present the fitted input function (black lines) and the distribution of feature-contrast in the white Gaussian noise stimuli (grey histograms, Fig. 3a1). The same format is used in Figures 3b1, 3c1.

For the cell in Figure 3a, the estimated input nonlinearities for both filters are close to quadratic. We also show a selection of interpolated features that are associated with the locations in the feature subspace indicated by the adjacent green dots. The polar angle (with the x-axis) of each green dot is referred to as the feature-phase of the corresponding feature. It uniquely identifies the spatial form of the feature, independent of its contrast (refer also to Fig. 1d). The space of all spatial feature forms, neglecting contrast, is referred to as the feature spectrum. As the green dots all lay on the same magenta contour they generate the same spike rate (Fig. 3a1; see Supplementary Video 1 for movie). For this cell, the spectrum of interpolated features producing equal responses spans 2π radians of feature-phase due to the closed elliptical iso-response contours. Elliptical contours arise because the input nonlinearities are insensitive to image polarity for either primary features, thereby, given sufficient feature-contrast, all interpolated features lead to spiking responses, regardless of brightness polarity. The invariance to feature-phase did not depend on a specific choice of iso-response contour. That is when we normalised each contour to its minimum feature-contrast (= radial distance in the feature subspace, Fig. 1d), the contours showed little variation (Fig. 3a2). Therefore, the functional form of relative feature-contrast as a function of feature-phase was independent of spike rate.

The set of interpolated features that produce equal spike rates in the feature subspace reveals the selectivity or invariance of the cell to any feature characteristic. We characterised the interpolated features of the subspace in terms of their peak orientation, peak spatial frequency, and relative spatial phase. The example cell is invariant to spatial phase because it responds equally to interpolated features that are spanning the full 360° of spatial phase (thick magenta line, Fig. 3a3). Conversely, the progression in orientation and spatial frequency of the interpolated features on the iso-response contour spanned a small fraction of the possible range (thick green and orange lines, Figs. 3a3). Therefore, this cell is selective for orientation and spatial frequency but is largely spatial phase and brightness polarity invariant, consistent with the description provided by the Energy Model (Adelson and Bergen 1985; Emerson et al. 1992).

We also encountered many 2-filter cells whose iso-response contours in their feature subspace were not elliptical (Fig. 3b1). These cells were typically invariant to only a limited range of feature-phases because their iso-response contours were open (Fig. 3b1, 3b2), i.e. they passed through only part of the circle of feature-phase, independent of the spike rate. For the cell in Figure 3b spatial phase spanned only 210° (thick magenta line, Fig. 3b3), largely due to the one-sided input functions (black lines in plots adjacent to axes, Fig. 3b1). The primary features for this cell differed in spatial phase by 113°, in peak orientation by 13° and spatial frequency by 0.09 cpd. The relationships between the primary features result in noteworthy variations in the characteristics of the interpolated features (insets in Fig. 3b1; see Supplementary Video 2 for movie). For example, while this cell had limited invariance to spatial phase, it was invariant to relatively large perturbations in orientation and spatial frequency, i.e. ranges of 22.5° and 0.18 cpd, respectively (thick lines, Fig. 3b3). Moreover, the cell remained sensitive to feature polarity: it responded to only positive feature-contrast for both primary features (top right, Fig. 3b1).

Figure 3c1 shows the responses of a cell with one filter and an odd-symmetric input function (i.e. near-linear input function). It is difficult to compare the feature subspace of such a cell with that of cells with two filters. Therefore, for visual presentation we show the response in a 2-dimensional feature subspace spanned by the primary feature of its RF filter and an additional, dummy feature that does not influence responses (i.e. it is assumed to be orthogonal to the primary feature via inner product). The choice of the dummy feature does not affect the shape of the feature-contrast response function, which is characterised by linear, vertical iso-response contours (Fig. 3c1). This represents the extreme degree of openness in iso-response contours in our population, arising when the cell exhibits no response dependence along the dummy stimulus dimension. Cells that have slight response dependence on a primary feature approach this limit.

As the iso-response contours of the cells in Figures 3b1 and 3c1 are open, responses to interpolated features at either end of their contours could only occur if feature-contrast was increased significantly. Therefore, each cell’s selectivity and invariance are reliant on the range of feature-contrast considered. To quantify the degree of response invariance within the feature subspace, we defined a parameter called the feature-phase bandwidth (Fig. 3b2 and 3c2; see ***Methods***). This is the range of feature-phases capable of driving each cell at a fixed spike rate with less than twice the feature-contrast required by the optimum feature (Fig. 3b2 and 3c2). The optimum feature at each spike rate was determined as the interpolated feature requiring the minimum feature-contrast to drive that spike rate. For most cells, including the three example cells in this subsection, this procedure depended little on the choice of spike rate level because the shape of the iso-response contours was largely independent of spike rate (except close to spontaneous rate) so that the contours were approximately scaled (i.e. magnified) versions of each other. In two dimensions the feature-phase bandwidth has a maximum of 2π radians. The cell in Figure 3a attains this maximum value, indicating invariance across the full spectrum of features in the feature subspace. The feature-phase bandwidth of the cell in Figure 3b was 1.3π radians (Fig. 3b2), indicating invariance across a partial range of the feature spectrum. For cells with a single filter the feature-phase bandwidth can be calculated analytically: it is always 2π/3 radians for a 2-dimensional feature subspace and corresponds to the most limited degree of invariance across the feature spectrum we observed (Fig. 3c2). The feature-phase bandwidth is used in ***Part 4*** of the ***Results*** to characterise the feature-contrast response function and again in ***Part 5*** to quantify the overall selectivity and invariance of cells to feature characteristics.

The examples given in Figure 3 show that the selectivity and invariance of a cell’s response to different feature characteristics is a consequence of two independent factors: (1) the feature spectrum resulting from the cell’s primary features, which determines the potential range of interpolated feature characteristics to which the cell is sensitive; and (2) the feature-contrast response function of the cell that determines which part of this feature spectrum is sampled to give a response, as quantified by the feature-phase. The two factors will be analysed separately in ***Parts 3*** and ***4***, and the resulting combination of factors in ***Part 5***.

### Part 3. Population analysis of the feature spectrum

Here we quantify the characteristics of all interpolated features for each cell. The interpolated features exhibited considerable variability. For 68% of cells with at least two filters, 90% of the interpolated features were Gabor-like (Fig. 4a). For the remaining 32% the feature spectra also contained a substantial portion of blob-like features (Fig. 4b) or features that are difficult to classify into a single category, which we term “unclassified” (Figs. 4c).

We examine the diversity of Gabor-like features in each cell’s spectrum in terms of peak orientation, peak spatial frequency and spatial phase. For each of these characteristics we considered the values covered by features in the cell’s spectrum, referred to as the spectral range for that feature characteristic (Fig. 5a-c). The spectral range is a measure of the diversity of a feature characteristic in a cell’s spectrum.

**Figure 5.**
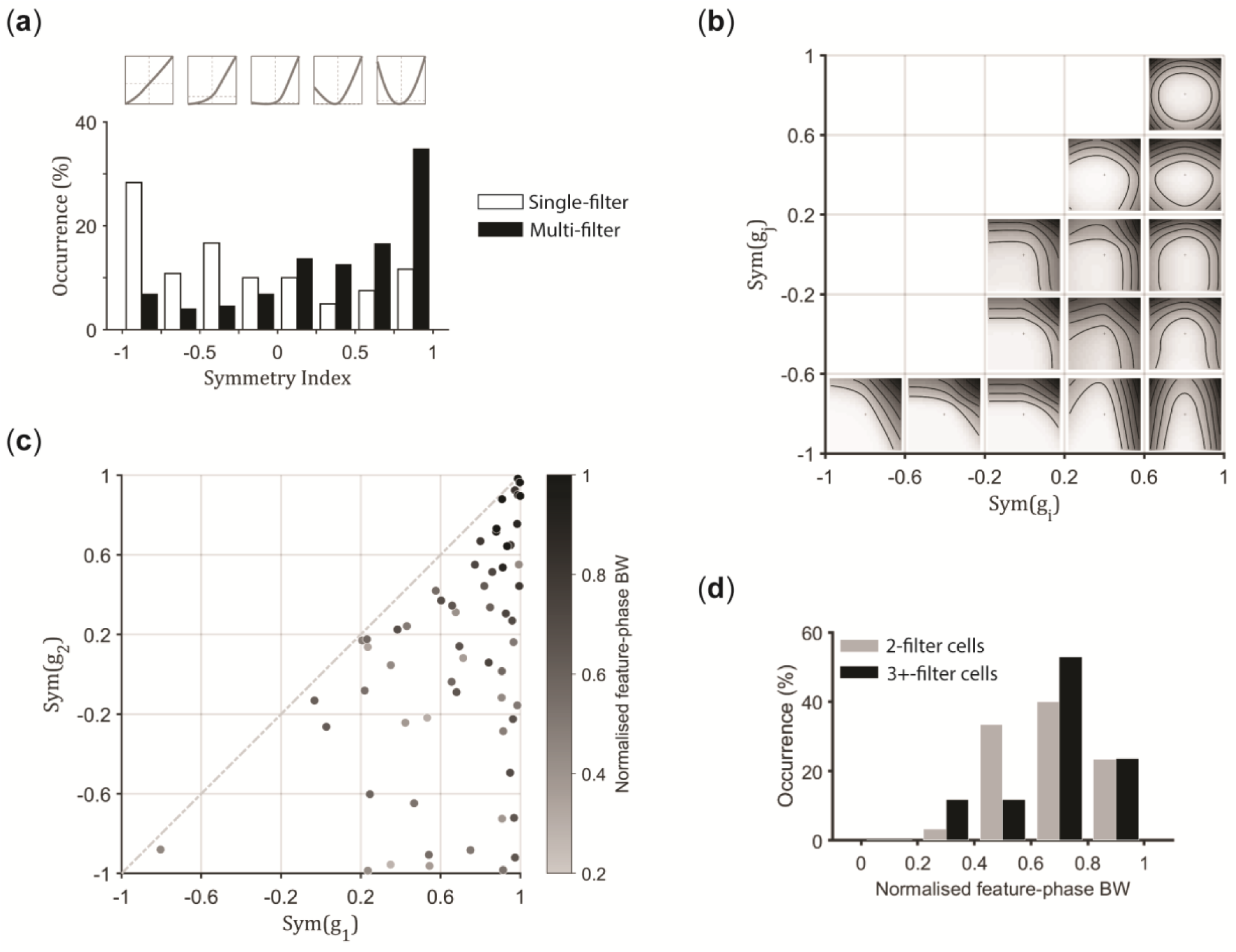
Diversity in the input functions of the V1 cells. (**a**) Distributions of the symmetry indices of the identified input functions for individual filters in our population of cortical cells, split into single-filter (white bars), versus multi-filter cells (black bars). Inset shows example input functions, in ascending order (from left to right) of symmetry indices. (**b**) Spectrum of 2D feature-contrast response functions along with the iso-response contours acquired by combining pairs of input functions with different forms indicated by the symmetry indices. (**c**) Scatter plot depicting the diversity in the combination of input functions, in terms of symmetry indices, found for cells that pool across two primary features. The grey level of each data point indicates the cell’s normalised feature-phase bandwidth (see colour bar). (**d**) Distributions of feature-phase bandwidth given for cells with 2 filters (light bars) versus cells with 3+ filters (dark bars). Sym; symmetry, BW; bandwidth.

Orientation showed the least variation within a cell’s feature spectrum relative to its maximum value of 180°: e.g. for cells with 2 filters, 93% had preferred orientations within 45° of each other (light dashed line, Fig. 5a). Cells with 3 or more (3+) filters exhibited wider variations in preferred orientations, with 65% having a spectral range >45° (dark dotted line, Fig. 5a). This greater diversity of orientation preferences may simply be because cells with 3+ filters have higher dimensional feature subspaces than those with 2 filters, resulting in a greater number of features. However, this did not stand for cells with >90% Gabor-like features, where all cells were limited to spectral ranges for orientations that were <45° (bottom right corner, Fig. 5d), regardless of whether they had 2 or 3+ filters (light squares versus dark circles, respectively). The cells with spectral ranges for orientation >45° mostly came from cells that had >10% non-Gabor-like features and 3+ filters, e.g. in Figure 5d 73% of cells showing an orientation range >45° had 3+ filters (dark circles) and all have >10% non-Gabor-like features.

Spatial frequency showed intermediate degrees of variation within a cell’s feature spectrum relative to its maximum value of 0.6 cpd: 52% of cells with 2 filters had spectral ranges <0.1 cpd (light dashed line, Fig. 5b). Cells with 3+ filters tended to have larger spectral ranges (dark dotted line, Fig. 5b). However, this did not appear to be limited to cells with >10% of non-Gabor like features in their spectra (Fig. 5e).

Spatial phase tended to show the greatest degree of variation within a cell’s feature spectrum relative to its maximum of 360°, with 75% attaining the maximum or near maximum spectral range (Fig. 5c). The difference in the spectral ranges for spatial phase between cells with 2 versus 3+ filters was not pronounced amongst cells that contained >90% of Gabor-like features (Fig. 5f). However, for cells with 2 filters that contained >10% of non-Gabor-like features in their spectra, the spectral range for spatial phase tended to diminish in proportion to the fraction of Gabor-like features, suggesting that the quantity of Gabor-like features in the spectra was insufficient to cover the full range of spatial phase in these cases (Fig. 5f) (note that the spatial phase, spatial frequency and orientation are only defined for Gabor-like features). In contrast, cells with 3+ filters typically attained the maximum spectral range of 360° for spatial phase regardless of the portion of Gabor-like features in their spectra (Fig. 5f).

### Part 4. Population analysis of input functions and feature-contrast response functions

The input functions of neurons varied from even-symmetric curves (invariant to feature-contrast polarity, e.g. black traces in subplots of Fig. 3a1 adjacent to the main plot’s axes), through one-sided functions (responding to one polarity of feature-contrast, Fig. 3b1), to odd-symmetric functions (excited by one polarity and inhibited by the opposite polarity, Fig. 3c1). The input functions were characterised by a symmetry index around zero feature-contrast. For cells with single filters the distribution peaked on odd-symmetric functions (white bars, Fig. 6a; n=120). For multi-filter cells the input functions were biased towards even-symmetric (black bars, Fig. 6a; n=77). Nonetheless for many cells with multiple filters, the input functions departed from the even-symmetric form.

**Figure 6.**
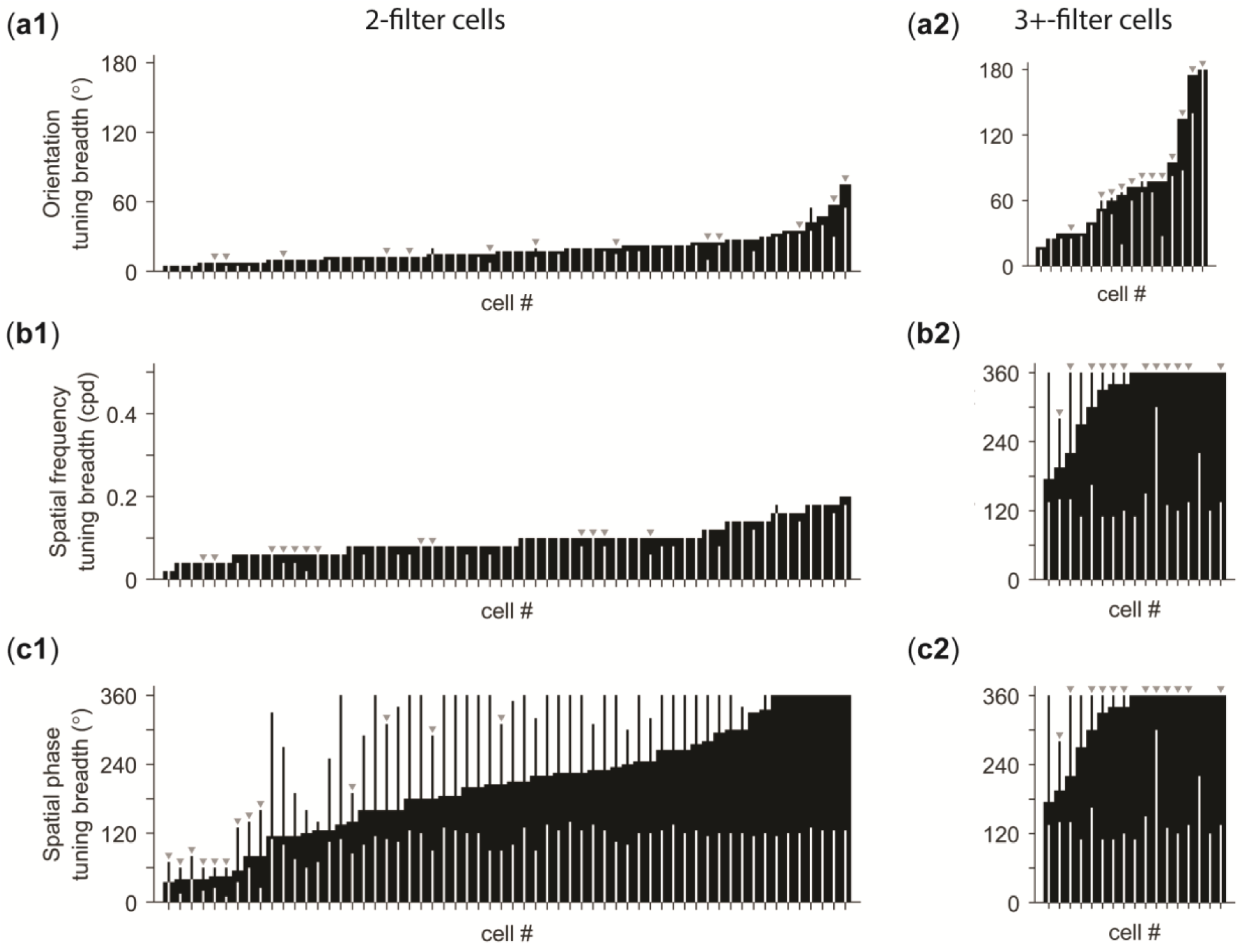
Selectivity or invariance for V1 cells to basic feature characteristics. Bar graphs present tuning breadths of cells with 2 filters (left column) versus cells with 3+ filters (right column), for (**a**) orientation, (**b**) spatial frequency, and (**c**) spatial phase. Each tick mark on the abscissa of each bar graph indicates a single cell. The length of each thick black bar indicates the tuning breadth for each cell. The length of thin black bars that occasionally rise above thick bars indicates the spectral range in feature characteristics. The length of white thin bars indicates the breadth if a linear coding scheme was applied for pooling (as in Fig. 3**c**). The light triangular symbols above the bars denote cells whose feature spectra contained <90% Gabor-like features.

When the input functions of filter pairs were combined via additive pooling and the spiking nonlinearity (as per the NIM, Fig. 1c), the variety of input functions led to a range of 2-dimensional iso-response contour shapes for the feature-contrast response functions in the cell population. Figure 6b shows example functions for cells with two filters. Cells in the top right of the plot had closed iso-response contours of circular or elliptical form that combined two even-symmetric input functions and responded to all values of feature-phase. Other cells had curved open iso-response contours that varied in shape from arches to more obliquely open forms that responded to limited ranges of feature-phase. These arose when at least one input function of a one-sided or odd-symmetric type was incorporated into the overall feature-contrast response function (Fig. 6b). In the bottom left of the plot the iso-response contours approach straight lines (seen in the linear model (e.g. Fig. 3c)), which occurs when at least one of the input functions is odd-symmetric. From a scatter plot of pairs of symmetry indices for input functions of cells pooling across two filters, the most frequent cell class had a pair of input functions with even-symmetric types (Fig. 6c). However, cells representing all other shapes are represented, revealing a diversity of input and feature-contrast response functions (Fig. 6b).

The portion of the feature spectrum encompassed by the feature-contrast response function was quantified by its feature-phase bandwidth. The points in Figure 6c have been shaded according to the normalised feature-phase bandwidth (between 0 and 1). This shows that cells with pairs of even-symmetric input functions are closely associated with the broadest feature-phase bandwidths (dark circles) corresponding to closed iso-response contours and those with at least one odd or one-sided input function have narrower bandwidths corresponding to open contours (light circles).

For cells with 3+ filters (n=17, ∼9%) the iso-response contours form a surface (or hypersurface) that can be classified in a similar way to the 2-dimensional examples, i.e. closed, or open and curved, or open and approximately linear (i.e. planes). In these higher dimensional cases, the feature-phase bandwidth can be generalised conceptually as the (solid) angular fraction of the feature spectrum enclosed by the contours. In analogy to the 2-dimensional case, we defined this as the fraction of the “surface area” of a hypersphere centred on the origin, for which the relative feature-contrast required to drive the cell was less than twice the minimum value. For an ideal higher-dimensional model with closed hyperspherical contours, this normalised feature-phase bandwidth is 1 (note the surface area of a hypersphere varies with dimension, so normalising by this area allows comparison across dimensions). For the ideal linear model with hyperplane contours, the normalised feature-phase bandwidth decreases with the dimension of the feature subspace from 1/3 ≈ 0.33 in 2 dimensions to 5/32 ≈ 0.16 in 5 dimensions. Five significant filters formed the highest dimensional feature subspace in our population of cells. Across the population, cells with 2 and with 3+ filters both had a wide-distribution of feature-phase bandwidths spanning these limits (Fig. 6d).

An important difference between cells with open versus closed contours was that open-contour cells responded to only part of the features in the feature spectrum. As a consequence, for open-contour cells the feature that drove the cell with minimum feature-contrast would not typically drive the cell when the brightness polarity was reversed (e.g. Fig. 3c). Conversely, the responses of closed-contour cells were approximately invariant to the brightness polarity of the feature (e.g. Fig. 3a).

### Part 5. Population analysis of overall response selectivity and invariance

The previous sections have shown the full spectral range of interpolated features (***Part 3***) and the types of input functions, that lead to the final output in the form of the feature-contrast response function characterised by its feature-phase bandwidth (***Part 4***). We now combine both stages of the model to establish the spectral range after the steps of applying the input functions, pooling and spiking nonlinearity. These steps can restrict the spectral range according to the cell’s feature-phase bandwidth because the cell does not respond over the full spectrum of features, as shown in Figure 3b. Therefore, it is necessary to use a different term for the restricted spectral range of a feature characteristic, which we call the characteristic’s tuning breadth. The tuning breadth of a feature characteristic only includes interpolated features into the restricted spectral range if their feature-contrasts are less than two times the minimum feature-contrast required to maintain the response, as defined for the feature-phase bandwidth. For example, Figure 3b2 shows the feature-phase bandwidth of an example cell and Figure 3b3 shows the corresponding tuning breadths for orientation, spatial frequency and spatial phase (projection of thick lines onto the vertical axis) as restricted from the full spectral ranges (projection of the extended curves by thin lines onto the vertical axis).

We present the tuning breadths estimated from multi-filter cells along with the theoretical tuning breadths that would be obtained if the cells had access only to linear or quadratic-like input functions. The tuning breadth using a quadratic-like nonlinearity is equivalent to the full spectral range because it gives circular iso-response contours with access to the full feature-phase bandwidth (e.g. cell in Fig. 3a). At the opposite end of the continuum is linear spatial processing, which has the narrowest feature-phase bandwidth (e.g. cell in Fig. 3c).

Figure 7 shows the diversity in tuning breadths for orientation, spatial frequency and spatial phase across the population of 77 cells with 2 or more filters. The left-hand column shows cells with 2 filters, whereas the right-hand column shows the cells with 3+ filters. The grey triangular symbols above the bar-plots highlight cells (n=52) that had <90% Gabor-like features in their feature spectrum, which over-emphasised blob-like or unclassified features. In these histograms, thick bars show the tuning breadths obtained from the cells. Thin black bars show the tuning breadth if the cell spectrum is sampled using a combination of quadratic-like input functions. Thin white bars show the tuning breadth if the linear model, aligned to the interpolated feature to which the cell responds at minimum feature-contrast, samples the cell spectrum. These quantities form upper and lower bounds to the cell’s tuning breadth, respectively, as they correspond to maximal and minimal feature-phase bandwidths.

**Figure 7.**
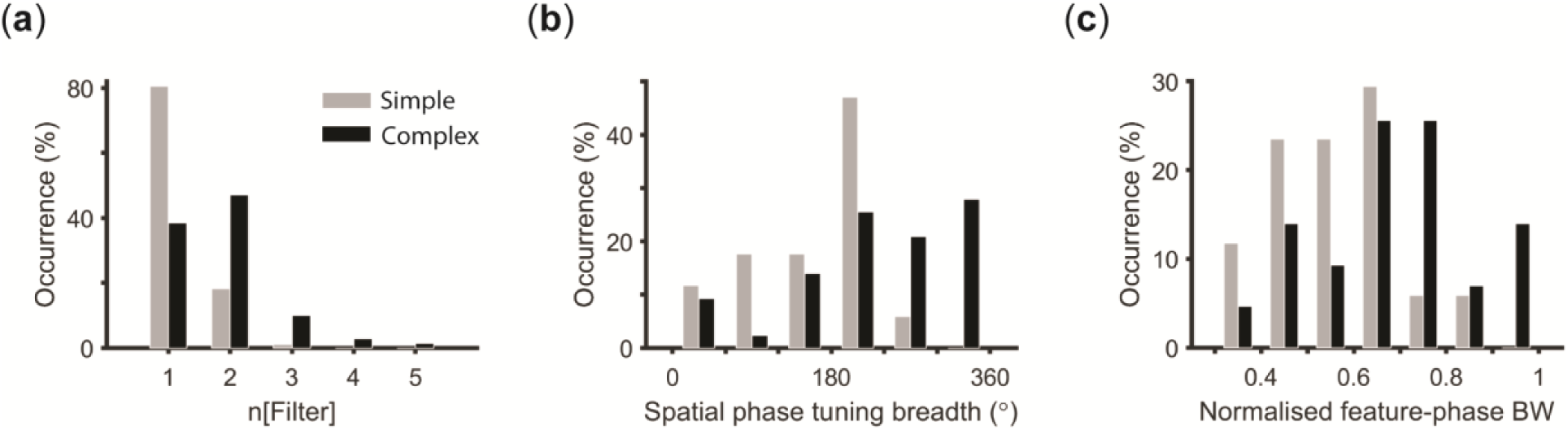
Correspondence to classical simple and complex cells. The bar graphs show distributions of (**a**) number of RF filters for each cell according to NIM characterisation, (**b**) the tuning breadth in spatial phase, and (**c**) the normalised feature-phase bandwidth (BW), for our population of V1 cells, split into simple (light bars) and complex (dark bars) cells according to their classification by response modulation ratios to sinusoidal drifting gratings.

Most cells with 2 filters showed a high degree of selectivity for feature orientation (95% had orientation tuning breadth <45°; thick bars, Fig. 7a1). However, the subpopulation of cells with 3+ filters exhibited invariance to larger perturbations in orientation particularly if they also had >10% non-Gabor-like features in their feature spectra (92% had tuning breadth >45°; grey triangles, Fig. 7a2). For most cells with 2 filters, their nonlinear feature-contrast response functions (thick black bars) did not sample a greater range of orientations in the feature spectrum than the linear model (thin white bars, Fig. 7a1), but in nearly all cases this still corresponded to the full spectral range allowed by the model with quadratic-like input functions (thin black bars rising occasionally above the thick black bars, Fig. 7a1). This is because values of orientation are often repeated cyclically around the feature spectrum so that the full range of orientations can be sampled from just a portion of the spectrum (e.g. Fig. 3b3). However, many 3+-filter cells with >10% non-Gabor-like features sampled a wider range of orientations than expected from the linear model, offering significantly greater invariance to orientation (thick black versus thin white bars with grey overhead triangles, Fig. 7a2).

Most multi-feature cells showed a high degree of selectivity for spatial frequency (61% tuning breadth <0.1 cpd, maximum 0.6 cpd). Most cells also had a spatial frequency tuning breadth equal to the maximum allowed by the model with quadratic-like input functions, i.e. the thin black bars rarely rise above the thick black bars (Fig. 7b1, b2). The cells with the greatest spatial frequency tuning breadths had 3+ filters (Fig. 7b2). Some of these cells also exhibited a modest advantage in broadening their spatial frequency tuning breadths as a result of the non-linear sampling of their response functions, i.e. the thick black bars are taller than the thin white bars (Fig. 7b1, b2).

Surprisingly, over the whole population of cells with 2 filters, only 12% had spatial phase tuning breadths that covered 360° (thick bars, Fig. 7c1). The subpopulation with 3+ filters mostly had spatial phase tuning breadths close to 360° (Fig. 7c2). In the broader population, 79% of multi-feature cells had spatial phase tuning breadths between the upper 360° limit and the lower limit of the linear model. They exhibited only partial spatial phase invariance but more than is possible from linear processing (thick black bars versus thin white bars, Fig. 7c1, 7c2). For most of these cells, this partial level of spatial phase invariance was entirely due to their nonlinear feature-contrast response functions, which only partly sampled the underlying feature spectrum. Hence, partial phase invariance was not because the spectral range of spatial phases in the cell’s feature spectrum was <360°. Those cells that had a limited spectral range for spatial phase were largely contained in the subpopulation of cells pooling 2 features and a high proportion of non-Gabor-like features (grey triangles, Fig. 7c1). A significant portion of these cells had very low spatial phase tuning breadths (<120°) because many features in their feature spectra did not have a defined spatial phase.

## DISCUSSION

We have used a general class of neural model (the NIM) to estimate the spatial features to which cells in V1 are sensitive and the nonlinear processing they use to integrate combinations of those spatial features in their responses. The approach was applied to multielectrode recordings of single units in anaesthetised cats in response to spatially white Gaussian noise. The estimated models allowed us to describe the nonlinear processing that V1 neurons employ to achieve tuning to various feature characteristics along a continuum from selective to invariant. The main feature characteristics we considered were orientation, spatial frequency and spatial phase for Gabor-like features.

In the NIM, selectivity and invariance of a cell’s response to different feature characteristics is a consequence of two independent factors: (1) the feature spectrum, which determines the potential range of feature characteristics to which a cell is sensitive; and (2) the feature-contrast response function that determines which part of the feature spectrum is sampled to give a response. The two factors were analysed separately in ***Parts 3*** and ***4*** of the Results, respectively, and their final combination in ***Part 5***.

For cells with 2 filters, most were relatively selective for feature orientation and spatial frequency (95% had orientation tuning breadth ≤45°, maximum 180°; 78% spatial frequency tuning breadth ≤0.1 cpd, maximum 0.6 cpd). By contrast, these 2-filter cells exhibited a wide variety of tuning to spatial phase, from highly selective to completely invariant (i.e. spatial phase tuning breadths from <60° up to the maximum of 360°). The role of the feature-contrast response function in determining the tuning breadths of these 2-filter cells also differed between feature characteristics. For orientation and spatial frequency, the cell’s feature-contrast response function was always sufficient to sample the full spectral range and played little role in determining their tuning breadths. For spatial phase it played a critical role in sampling a limited portion of the spectral range, reducing the range from a value that for most cells was the maximal 360°. Therefore, the tuning breadth for spatial phase was largely a reflection of the feature-phase bandwidth of the cell, which quantifies the portion of the feature spectrum sampled by the feature-contrast response function.

For cells with 3+ filters, tuning breadths were typically higher than for cells with 2 filters: 65% had orientation tuning breadth >45° versus 5% for 2-filter cells; 100% had spatial frequency tuning breadth >0.1 cpd versus 22% for 2-filter cells; 71% had spatial phase tuning breadth >300° versus 15% for 2-filter cells. While this may be interpreted as showing that cells with 3+ filters typically exhibited greater invariance than those with just 2 filters, some care needs to be taken in interpreting this result as there are at least two alternative interpretations. First, it could be that cells with 3+ filters have larger tuning breadths than those with 2 filters, simply because we considered their response to a greater range and quantity of features as a consequence of the higher dimensionality of their feature subspaces. Consistent with this, the spectral ranges for orientation and spatial frequency were mostly higher across the population of cells with 3+ filters compared to those with 2 filters (Fig. 5), indicating the feature subspace contains features with a greater diversity of these two characteristics. However, these interpolated features are not arbitrary but instead are particular to each cell and constitute the subspace of all possible features to which the cell is sensitive. Thus, a second interpretation is that the greater diversity of feature characteristics in the spectra of cells with 3+ filters may be because the system selected these particular features in a particular cell’s spectrum to enhance selectivity or invariance to some of their characteristics.

Evidence regarding these alternatives comes from considering the tuning breadths for the linear model (white bars, Fig. 7), which effectively reduces the number of filters in the model to just 1 while still estimating the tuning breadths in the original, larger dimensional feature subspace. The first observation is that the tuning breadths for the linear model are typically larger for cells with 3+ filters than those with 2 filters. This indicates that if we reduce the effective dimensionality of the model to a single filter then the dimensionality of the feature subspace used to estimate the tuning breadth will affect its value. The second observation is that for cells with 3+ filters, nonlinear processing due to the cell’s feature-contrast response function is contributing to the greater tuning breadths for orientation and spatial frequency compared to the linear model. This is not the case for cells with only 2 filters. Together, these observations indicate that nonlinear processing moderately enhances invariance to orientation and spatial frequency in cells with 3+ filters but not those with 2 filters.

### Comparison to Standard Models

An important difference between our study and previous work is that we have chosen to characterise the properties of all features in the feature subspace of the cell, instead of just the primary features corresponding to the model filters. Previous studies used STC (or related methods that analyse eigenvectors of a symmetric matrix), which forces the filters to be orthogonal. As emphasised by several authors (Rust et al. 2005; Touryan et al. 2005; Kaardal et al. 2013), such orthogonal filters should not be considered to have any anatomical interpretation as presynaptic inputs but rather to embody an arbitrary basis for representing the cell’s feature subspace. While the NIM was originally devised to find a functional basis that might allow anatomical interpretation, we did not take this approach. Rather, we preferred to analyse all features in the subspace, as they correspond to different choices of basis to represent the subspace.

This study is the first to provide a quantitative analysis at the population level of the feature-contrast response function. As a result, we found that the input functions of complex cells are not generally quadratic-like and the feature-contrast response functions of these cells do not generally have closed iso-response contours (Touryan et al. 2002, 2005; Rust et al. 2005; Chen et al. 2007). An important difference between our study and previous work is that we have estimated the nonlinear feature-contrast response functions using maximum likelihood estimation of the NIM, while previous studies used STC. NIM allowed us to estimate a general form of the feature-contrast response function, consisting of arbitrary input functions, additively pooled and then passed through a spike nonlinearity that captures threshold, gain, convexity and spontaneous rate. This allows the feature-contrast response function to be estimated in relatively high dimensional feature spaces (>2) and without assumptions about the orthogonality of filters. STC methods struggle to estimate the complete feature-contrast response function for cells with more than 2 filter dimensions because they use a method for which the number of parameters scales exponentially with dimension (compared to linearly for the NIM). Consequently, for cells with 3+ filters they have frequently estimated only 1 or 2-dimensional response functions by marginalising (i.e. averaging) over the remaining dimensions. Alternatively, they have assumed the response function can be parameterised by the sum of linear and quadratic functions. In general, neither approach is ideal.

We found that the number of RF filters varied across the population from 1 to 5 (Fig. 2b). The standard models for cells in V1 contain 1 filter for simple cells (i.e. the Linear Model) and 2 for complex cells (i.e. the Energy Model). Using a classification of simple and complex cells based on the modulation ratio to drifting gratings (F1/F0 ratio), many simple and complex cells in our population accorded with these filter numbers (81% of simple cells had 1 filter, 47% of complex cells had 2 filters, Fig. 8a). However, a substantial portion differed, with some simple cells having 2 filters, and some complex cells having 1 or as many as 5 filters.

For complex cells, the Energy Model emphasises spatial phase invariance which is achieved by using two filters that are Gabor-like, with matched tuning for orientation and spatial frequency, but being 90° spatial phase shifted (Adelson and Bergen 1985; Emerson et al. 1992). The Energy Model also requires that the filter outputs are combined by a sum of input functions that are approximately quadratic. Only a minority of complex cells (defined using the F1/F0 ratio) in our population approximated this description. In general, we found greater variety than suggested by the Energy Model in the combinations of filters employed by cells, their input and feature-contrast response functions and the forms of invariance they exhibited. The most significant departure was that we frequently found complex cells that exhibited only partial spatial phase invariance, which, at the lower end of the range, was similar to that of simple cells (Fig. 8b, spatial phase tuning breadths: complex cells 40° to 360°; simple cells 35° to 265°). Complex cells with partial spatial phase invariance were often selective for the polarity of the feature that elicits a given response at the lowest contrast, whereas the Energy Model imposes polarity invariance. The limited spatial phase invariance was primarily a consequence of limited sampling of the feature spectrum by the feature-contrast response function. This was quantified by the normalised feature-phase bandwidth, which varied from 0.3 to 1, overlapping considerably at the lower end of this range with the simple cell distribution (Fig. 8c). This variety for complex cells reflects the variety of their input functions, which were not just quadratic-like, but also one-sided and odd-symmetric. Finally, filters were not always well matched in orientation and spatial frequency, so that the spectrum of features to which the cell had approximately invariant response could exhibit perturbations in the orientation or spatial frequency of the features. Cells with 3+ filters were almost entirely complex cells in our population. As noted above, these cells moderately enhanced invariance to feature orientation and spatial frequency by using nonlinear processing.

A key concept distinguishing simple and complex cells, implicit in the conceptual models of Hubel and Wiesel (1962), is that of linear versus nonlinear spatial summation. This categorical distinction was made based on qualitative RF mapping techniques using spots, slits and bars as stimuli. Later work introduced sinusoidal grating stimuli, in part because sinusoids play a special role in linear systems, as a sinusoidal signal is preserved by linear processing, including its frequency, with changes in only the amplitude or phase of the input (Movshon et al. 1978a, 1978b; Dean and Tolhurst 1983). The standard Linear Model of simple cells provides a phenomenological account of their modulated response to drifting gratings that preserves the temporal frequency. For complex cells, the response to drifting sinusoidal gratings is less modulated than for simple cells, as quantified by the modulation ratio, provided the orientation, spatial and temporal frequencies are optimal in driving the cell. The standard Energy Model provides an explanation for this spatial phase invariance at a computational level. The bimodal distribution of the modulation ratio across the V1 population has supported arguments in favour of a dichotomous classification of simple and complex cells based on their response to optimal drifting gratings, supporting the importance of the two standard models (Skottun et al. 1991). However, this dichotomy has been questioned (Chance et al. 1999; Mechler and Ringach 2002).

For non-optimal grating parameters, modulation of complex cell responses to drifting gratings is frequently observed, e.g. below a cell’s peak spatial frequency. The Energy Model does not account for this modulation. A complete characterisation of a linear response using sinusoidal gratings requires measurements of responses across a sufficiently broad range of gratings, covering different orientations, spatial and temporal frequencies and spatial phases, and not just the peak values. Further, complete characterisation of a nonlinear response requires measuring responses to superpositions of all these sinusoids. Other stimulus sets, such as white noise and natural images, are also sufficiently rich to allow complete characterisation of nonlinear responses. These kinds of stimuli have been used to estimate highly quantitative models of simple and complex cells that aim to completely characterise the cell’s responses by using computational techniques such as spike triggered average and covariance techniques (Touryan et al. 2002, 2005; Rust et al. 2005; Chen et al. 2007), or Wiener-Volterra analysis (Fournier et al. 2014).

Early studies employing these stimulus sets and techniques estimated models that were largely consistent with standard models for simple and complex cells (Touryan et al. 2002, 2005). For example, Touryan et al. (2002) found that 78% of the complex cells had two significant filters that were quadrature pairs. Their measured contrast-response functions resembled quadratic nonlinearities with contrast polarity invariance, as expected of the Energy Model. However, later studies have estimated models with more filters than described in the standard models for simple and complex cells (Rust et al. 2005; Chen et al. 2007; Fournier et al. 2014). For example, Rust et al. (2005) found multiple filters for all the simple cells in their population. Similarly, for complex cells they found more than two significant RF filters - both excitatory and suppressive. In general, these later studies have emphasised the diversity of filters in terms of their relative orientation, spatial frequency and excitatory or suppressive nature (Chen et al. 2007; Fournier et al. 2014). They found that both simple and complex cells incorporate forms of nonlinear spatial summation that are not accounted for in the standard models for each cell type.

Our results generally agree with the later studies. The discrepancy between our findings and those of Touryan et al. (2005) could arise for a variety of reasons. The different model estimation studies have varied in a number of ways, including the type of stimuli (random bars, random squares, natural images), the species (cat, macaque), the state of consciousness (awake, anaesthetised), the type of response recorded (spikes or intracellular membrane potential), the quantity of spikes, and finally the model form and computational method for model estimation (STA/STC, Wiener-Volterra analysis, NIM with maximum likelihood). The first three differences cannot alone account for the discrepancies between Touryan et al. and later studies, as cats, anaesthesia and STA/STC were common features. In fact, Touryan et al. (2005) did report a portion of cells with 3+ filters in cat V1, similar to those found in our study. However, these extra filters were not considered further as it proved impossible to confirm their status because of the known susceptibility of STC analysis to generate artefacts (Paninski 2003; Sharpee et al. 2004; Schwartz et al. 2006), unlike the maximum likelihood method used here. Nonetheless, by reviewing all these model estimation studies, including the present investigation, it appears that there are differences in the number of estimated filters: (1) extracellular recording generates more filters in primates than in cats (Rust et al. 2005; Chen et al. 2007) and intracellular recording generates more filters in cats than does extracellular recording (Fournier et al. 2014). It should also be noted that the use of natural scenes over white noise may result in more filters.

In summary, RF mapping studies using rich stimulus sets, including this study, have revealed that a population of simple and complex cells use more elaborate nonlinear spatial summation than accounted for by standard models. Our results indicate that these elaborations have a significant impact on the relative selectivity and invariance of cells in primary visual cortex.

### Possible explanations of the observed selectivity and invariance

The degree of invariance in our V1 population was greatest for spatial phase, considerably less for spatial frequency and less again for orientation, relative to the maximal range for each of these characteristics. In the context of invariant object recognition in the ventral visual pathway, all three forms of invariance are important as they relate to translation, rotation and scale invariance in object recognition. There are several (non-exclusive) explanations for the differing degree of invariance we observed in V1. A first, teleological, explanation is that this may reflect the differing degrees of variability of these characteristics in Gabor-like features in natural scenes at the spatially local level relevant to V1 receptive field sizes. Objects deemed to be the same or similar may generally have only minor perturbations in the relative orientation or spatial scale of their constituent features at this spatially local level. Differences in the spatial positions of these features may be more considerable at the local scale.

A second, aetiological, reason favouring spatial phase invariance over other forms of invariance at the early cortical stages of processing concerns the dimensionality of the feature subspace. The spatial phase of a Gabor feature is an unusual feature characteristic, because the full 360° of spatial phase can be represented in a feature subspace of just two dimensions using a pair of Gabor features differing in spatial phase by 90° but otherwise identical. This is not the case for orientation or spatial frequency: the sum of two Gabor features with differing orientation or spatial frequency is not another Gabor feature, but will be approximately Gabor only when the difference in orientation or spatial frequency of the two features is not too large (i.e. comparable to, or smaller than the feature’s bandwidth for orientation or spatial frequency, respectively). Thus for 2- or 3-dimensional feature subspaces it is only possible to achieve small perturbations in the orientation or spatial frequency of the interpolated features in continuous fashion across the subspace. It may be difficult for neurons in earlier cortical stages of processing, like V1, to acquire sensitivity to features in subspaces with higher dimension than 2 or 3.

A third, aetiological, reason favouring spatial phase invariance over orientation or spatial frequency concerns the scale of the corresponding topographic maps in cortex. Maps for each of these characteristics have been demonstrated in cat primary visual cortex, but the spatial scale and

refinement (orderliness) of the maps differs, with orientation showing the most refined and broadest scale of organisation, followed by spatial frequency and then spatial phase (Nauhaus et al. 2012; Wang et al. 2015; Alonso 2016; Kremkow et al. 2016; Nauhaus et al. 2016). In the NIM, invariance arises due to the pooling of inputs across filters that have different feature characteristics. If we consider this pooling to correspond to the presynaptic input to the cell, then spatially local pooling would access the greatest diversity of presynaptic inputs for the characteristic of spatial phase, due to its fine spatial scale and greater randomness of organisation, followed by spatial frequency and then orientation.

## Acknowledgements

This work was supported by the Australian Research Council Centre of Excellence for Integrative Brain Function (CE140100007), the National Health and Medical Research Council (GNT1106390), and Lions Club of Victoria. AA, HM and MI would like to thank Anthony Burkitt and Colin Clifford for their valuable comments on the manuscript.

